# Supervised machine learning identifies impaired mitochondrial quality control in β cells with development of type 2 diabetes

**DOI:** 10.1101/2025.10.22.683335

**Authors:** Mirza Muhammad Fahd Qadir, Charles Dana, Paul Mauvais-Jarvis, Nazmul Haque, Theodore Dos Santos, Patrick E. MacDonald, Franck Mauvais-Jarvis

**Author notes:** Correspondence: MMFQ; FM-J.

## Abstract

In type 2 diabetes (T2D), molecular pathways driving β cell failure are difficult to resolve with standard single cell analysis. Here we developed an interpretable, supervised machine learning framework that couples sparse rule-based classification (SnakeClassifier), pathway constrained modelling (BlackSwanClassifier), and β cell mitochondrial fitness stratification (Kolmogorov-Arnold Neural Networks KANN), linking and integrating them into disease mechanisms in single cell RNA sequencing (scRNA-seq) from 52 human donors. SnakeClassifier trained on 50 genes accurately predicted T2D at single cell resolution, outperforming classical ensemble machine learning classifier models, and yielded donor level diabetes scores that correlated with chronic hyperglycemia. The clustering of β cell populations (β1-4) revealed a resilient non-diabetic (ND) β1 subtype characterized by preserved β cell identity genes and lower disease risk, whereas T2D β2-4 subtypes exhibited upregulation of genes involved in cellular and mitochondrial stress and suppression of genes promoting oxidative phosphorylation and insulin secretion. Mitophagy emerged as the dominant program linked to T2D and a mitophagy focused BlackSwanClassifier nominated *PINK1, BNIP3*, and *FUNDC1* as key regulators. *PINK1* was enriched in ND β1, decreased with T2D disease score and connected sex stratified mitophagy. We generated a KANN derived mitochondrial fitness index (MFI) integrating mitophagy, mitochondrial proteostasis, biogenesis and oxidative phosphorylation into a single interpretable score (R^2^ = 0.934 vs module-based mitochondria quality index), which identified mitophagy *PINK1, SQSTM1, PRKN* and *BNIP3* as top contributors to T2D progression. These transparent models unify prediction with T2D disease mechanism and identify the mitophagy receptor *PINK1* as a central determinant of β cell metabolic fitness

## Main

Type 2 diabetes (T2D) is a heterogeneous disease. This is reflected in the diversity of alterations in pancreatic islet cell populations, which remain incompletely understood^1^. Single cell RNA sequencing (scRNAseq) of human islets has enabled the construction of detailed atlases linking β cell transcriptional states to the progression of T2D^2-5^. For example, compared to non-diabetic individuals, female β cells of individuals with T2D exhibit suppressed genes involved in mitochondrial respiration, while male T2D β cells exhibit suppressed genes favoring insulin secretion, suggesting the existence of female-specific mitochondrial failure in the transition from normoglycemia to T2D^3^.

Despite advances in scRNAseq analysis workflows, resolving the complexity of β cell alterations in T2D remains challenging due to the transcriptional heterogeneity of islets from different donors, and confounding influences such as body weight, hyperglycemia and disease duration. These challenges underscore the need for novel computational strategies to mechanistically dissect the progression of T2D in human islet cells. Pathway aware machine learning (ML) approaches serve as the next frontier to interpret islet scRNAseq in the context of T2D complexity^5-7^.

Machine intelligence analysis of scRNAseq data enable trajectory inference^8,9^, dimensionality reduction^10,11^, data integration^12-15^ and disease modeling and prediction^16-18^. However, these models provide little mechanistic insight as they focus on noncausal statistical learning, not improving disease prediction. Recent advances in ML models have enabled cell annotation^19^, cell-cell interactions^20^, disease clustering^21^ and disease signatures^22^. Thus, by modeling the transcriptional heterogeneity of islet cells of T2D subjects using scRNAseq, ML offers the exciting possibility of predicting progression, treatment efficacy and adverse events, thus advancing precision medicine.

In this study, we leveraged ML to stratify, predict and mechanistically interpret β cell failure in the continuum of T2D. Using scRNAseq, as well as other experiments, we implement an interpretable ML rule-based classifier (SnakeClassifier) that learns sparse gene threshold rules, stratifies donors/sex and predicts a graded T2D β cell architecture. We identify a resilient β cell subtype in ND individuals. Using pathway-constrained ML (BlackSwanClassifier) and a Kolmogorov-Arnold Neural Network based mitochondrial function indexing, we identify impaired mitophagy as instrumental in β cell failure in T2D.

## Results

### Rule based interpretable learning of T2D β cell states using SnakeClassifier

In type 2 diabetes (T2D) impaired pancreatic islet cells genetic programs remain incompletely defined^1^. Resolving these programs at single-cell resolution requires a combination of models that overcome data sparsity and remain biologically interpretable. To address this, we developed and applied a Boolean rule-based classifier with logistic aggregation (SnakeClassifier) to define transcriptional differences between T2D and non-diabetic (ND) states in human pancreatic islets scRNAseq datasets **(Fig. 1a)**. Our framework is modality agonistic, the same objective functions that generate rule lists from RNA gene counts can also be applied to other feature sets such as sex, age, BMI. Runtime scales with the number of cells, avoiding the *O(nm*^*2*^) geometric computational complexity problem **(Supplementary Fig. 1a)**. SnakeClassifier and BlackSwanClassifier are interpretable by construction as each prediction is the logistic aggregation of a few *IF-THEN* gene threshold clauses. The trained model is sparse, demonstrating stable feature ranks (Kendall τ=0.48), and sign monotonicity (0.78) **(Supplementary Fig. 1b)**. We re-clustered β cells and utilized SnakeClassifier to explore T2D disease states **(Fig. 1b)**. We subjected clustered β cells to pathway over-representation analysis (ORA) to highlight biological mechanisms associated with disease progression and heterogeneity **(Fig. 1b)**. We also subjected selected upregulated pathways to a pathway constrained ML model BlackSwanClassifier enabling gene-ranking to identify critical biological regulators **(Fig. 1c)**. Functions such as mitochondrial quality rely on multiple pathways making their study based on gene expression challenging, termed as an integrated pathway cellular complexity problem. To address this challenge, we implemented Kolmogorov-Arnold Neural Networks on β cells clusters to model multiple integrated cellular pathways into a single additive function **(Fig. 1c)**. Collectively, this pipeline enables disease state prediction and interpretation of pathways in detail prioritizing candidate mechanisms associated with β cell failure.

**Figure 1.**
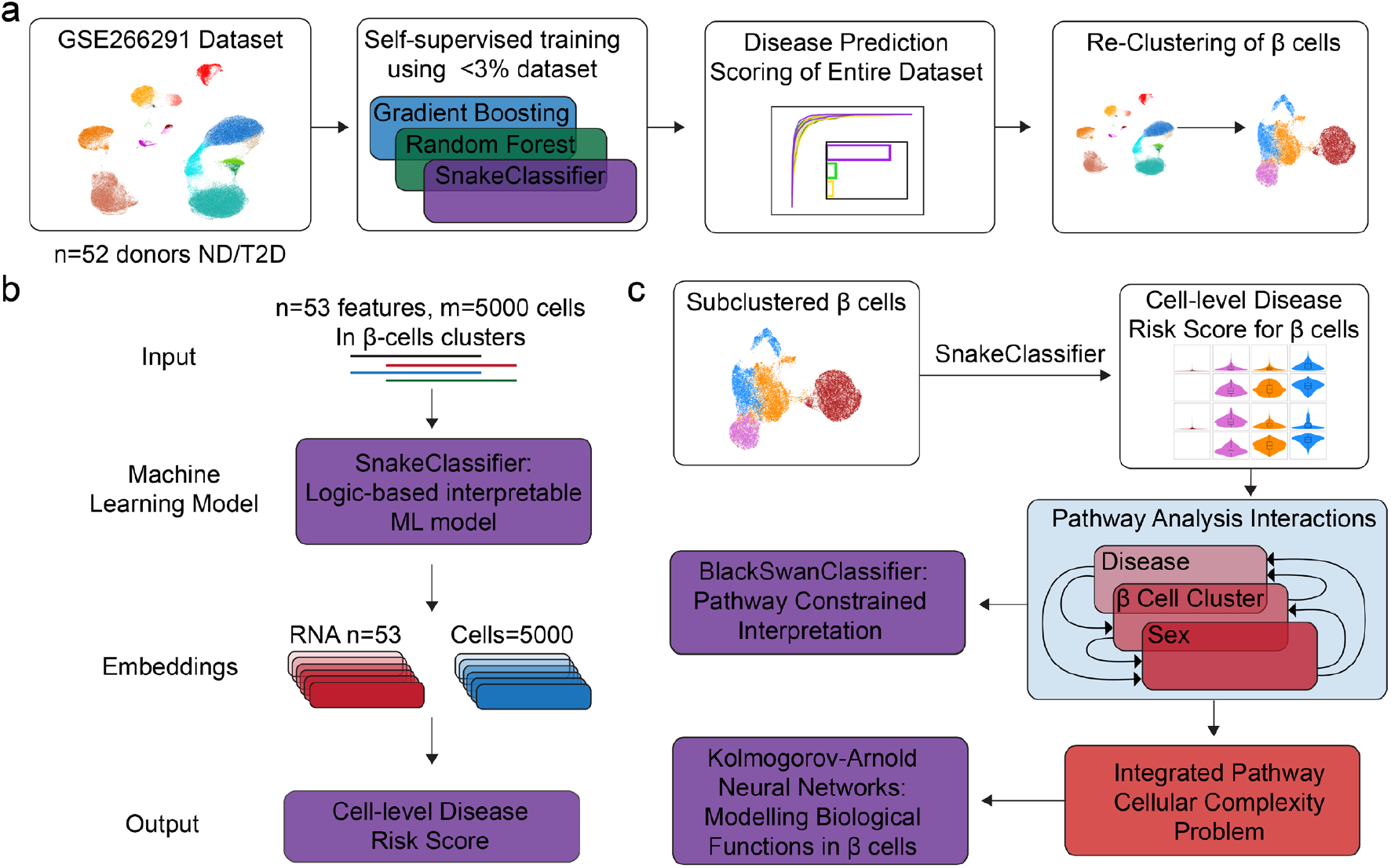
Schematic overview of cell-level disease state modeling and analysis. **a**, Overview of the analytical pipeline. Single-cell RNA-seq data from 52 human pancreatic islet donors (GSE266291, n=141,739 cells); non-diabetic [ND] and type 2 diabetic [T2D]) were used for model development. Less than 3% of the dataset was employed for self-supervised training of multiple machine learning classifiers, including Gradient Boosting, Random Forest, and the logic-based interpretable SnakeClassifier model. The trained models generated cell-level disease risk scores for the entire dataset, followed by re-clustering and annotation for analysis of cellular disease states. **b**, Schematic of the logic-based machine learning approach. Expression profiles for 53 features (including 50 genes) and 5,000 single cells were input into the SnakeClassifier model, generating cell and gene level embeddings that culminate in quantitative cell-level disease risk scores. **c**, Downstream analysis framework. Clustered β cells were subject to SnakeClassifier model to rank cells based on risk score. Clusters were subject to ORA using disease, cluster and sex as variables. The full dataset was classified by disease status, sex, and cell type. The SnakeClassifier model was applied to all cells (n=141,739), enabling disease scoring and the identification of disease and sex-specific states. β cell subsets with distinct disease states were isolated and subjected to pathway over-representation analysis (ORA) to reveal biological mechanisms associated with disease progression and heterogeneity. Pathways were subjected to BlackSwanClassifier for gene importance ranking. Complex cellular pathways modelling challenges 9termed integrated pathway cellular complexity problem) was interpreted using a Kolmogorov-Arnold Neural Network to model complex biological function in β cells.

### SnakeClassifier predicts T2D state from single cells with accuracy and clinical relevance

To predict which cells, exhibit a T2D phenotype, we implemented the logic-based interpretable ML model SnakeClassifier. Using 50 ranked genes, sex, ancestry and cell type as features (see methods for selection criteria), we trained SnakeClassifier on 5000 randomly selected annotated cells (3596 ND and 1404 T2D) and benchmarked against two other classical ensemble machine learning (ML) classifier models: Random Forest and Gradient Boosting, trained on the same features. All three models produced per-cell T2D risk scores for the complete annotated dataset (n=141,739)^3^. We evaluated performance using mean receiver operating characteristic (AUROC), which quantifies how consistently models rank true T2D cells above ND cells throughout sensitivity-specificity tradeoffs, and a weighted-winner statistics that identifies the superior model per comparison on the basis of effect size. All three models exhibited low false positive rate (**Fig. 2a)**. Calibration was stable: Youden’s J peaked at **J**^**\***^ = [0.78] at threshold **t**^**\***^ = [0.29], yielding sensitivity [0.91] and specificity [0.87] **(Fig. 2b** and **Supplementary Fig. 2a-b)**. Risk scores projected on UMAP maps separated ND from T2D states and showed higher risk score in T2D clusters for all cells **(Fig. 2c** and **Supplementary Fig. 2c-e)**. SnakeClassifier achieved the best single-cell T2D classification, with AUROC 0.958 in females and 0.948 in males, exceeding the self-supervised training of Gradient Boosting (F: 0.927, M: 0.884) and Random Forest (F: 0.948, M: 0.931 **Fig. 2d**). SnakeClassifier outperformed the two other models on most donors (weighted-winner, F: 80.4%; M: 95.1%; **Fig. 2e**). Predicted versus observed T2D gene composition per cell type and sex was tightly correlated (R = 0.93 in females and males; **Fig. 2f**). Out of the 50 genes selected by SnakeClassifier for disease state prediction (see methods for selection criteria), the top 10 ranked genes were increased in all T2D vs ND islet cell types **(Fig. 2g)**. We averaged the mean risk score for all cells in each donor to obtain the mean disease score for each donor. In that setting, chronic hyperglycemia (HbA1c) correlated with mean disease score (R = 0.68, p = 3.8 × 10^−8^; **Fig. 2h)**, indicating that our SnakeClassifier model of disease prediction exhibits both biological and clinical relevance.

**Figure 2.**
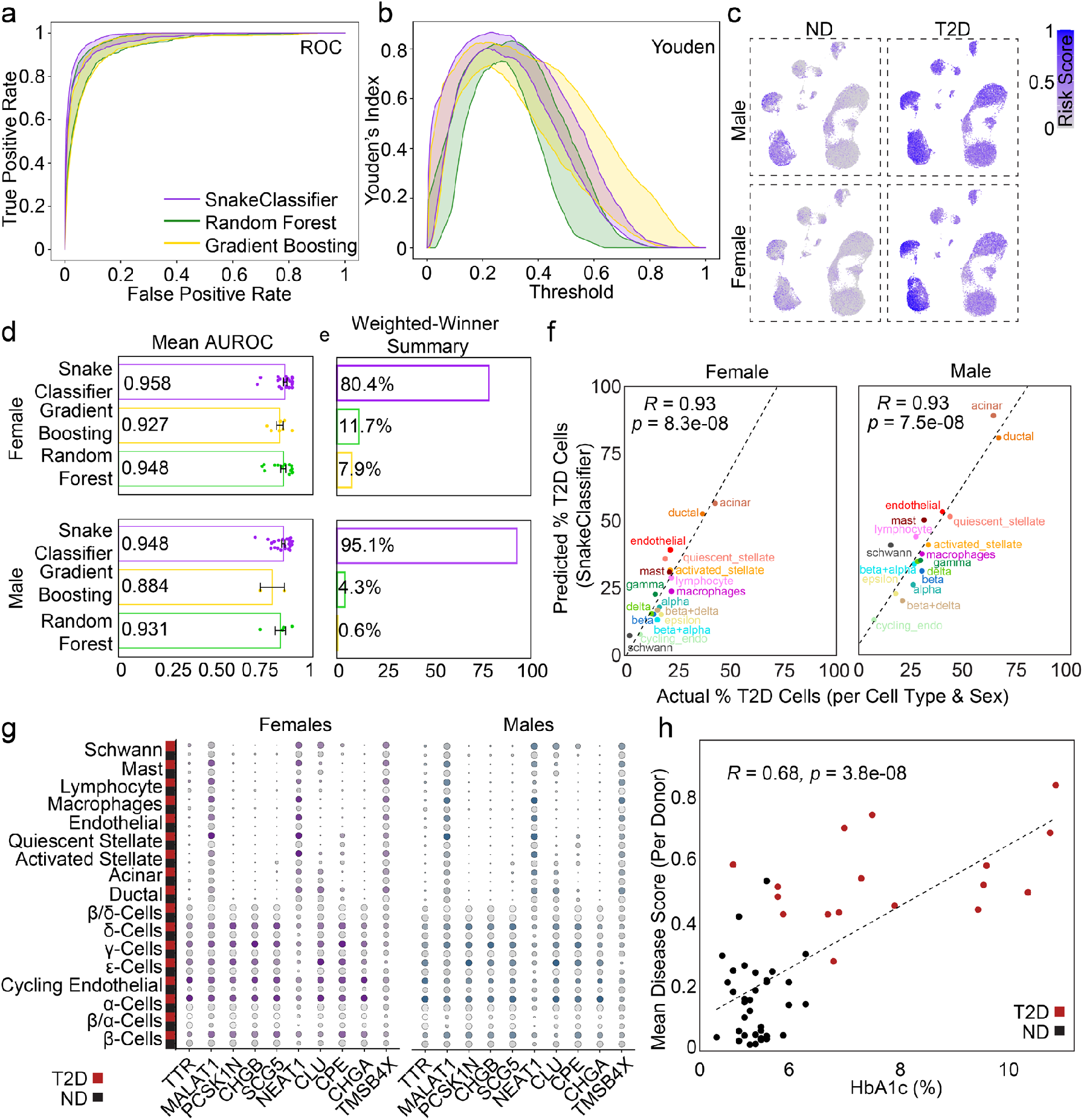
Model performance, cell-level prediction, and biological validation in human islets. **a**, Receiver operating characteristic (ROC) curves comparing Algorithm Snake, Random Forest, and Gradient Boosting classifiers for predicting T2D status at the single-cell level for all donors (n=52). **b**, Distribution of Youden’s index within prediction thresholds for each classifier, indicating optimal disease risk score cutoffs. **c**, UMAPs colored by predicted disease risk scores for each cell, stratified by sex (male, female) and disease status (ND, T2D), illustrating separation of T2D and ND cellular states. **d**, Model comparison summary showing mean area under the ROC curve (AUROC). **e**, Model comparison summary showing weighted-winner summary for each sex, highlighting the superior performance and consistency of SnakeClassifier within donors. **f**, Correlation between the predicted and actual percentage of T2D classified cells per cell type and sex (R = 0.93 for both females and males), demonstrating high model concordance with true disease status within diverse cell populations. **g**, Dot plot of cell-level disease risk scores for all major pancreatic islet cell types in male and female donors, of top 10 features, separated by disease status (ND, T2D). **h**, Scatter plot showing the correlation between mean disease risk score per donor and HbA1c (%) (R = 0.68, p = 3.8 × 10^−8^), supporting the biological relevance of the predicted cell-level disease scores.

### Cell type clustering reveals distinct β cell subtypes

The transcriptional diversity of β cells is reflected in glucose dependent insulin secretion^23-25^. We sub-clustered β cells from our previous study^3^ producing four β cell clusters (β1-4) **(Fig. 3a)** in which female T2D β2-4 and male T2D β3-4 exhibited enriched risk scores compared to ND β cells (**Fig. 3b)**. Note that male ND β2 clusters exhibited high disease score. Chronic hyperglycemia (HbA1c) correlated with increased disease scores for all β cell clusters except for β1 **(Fig. 3c)**. We quantified pathway activity in β cell clusters using Seurat’s module score function^26^, which integrates the expression of all genes in a pathway into a single composite signal called the mean module score. ND β1 showed the highest expression of β cell identity genes compared to ND β2-4 **(Fig. 3d)**, which was validated by pathway mean module scores **(Fig. 3e)** independent of sex. Mean module scores for β cell clusters were inversely correlated with insulin secretion and vesicle exocytosis in females, but not males **(Supplementary Fig 3a,b)**.

**Figure 3.**
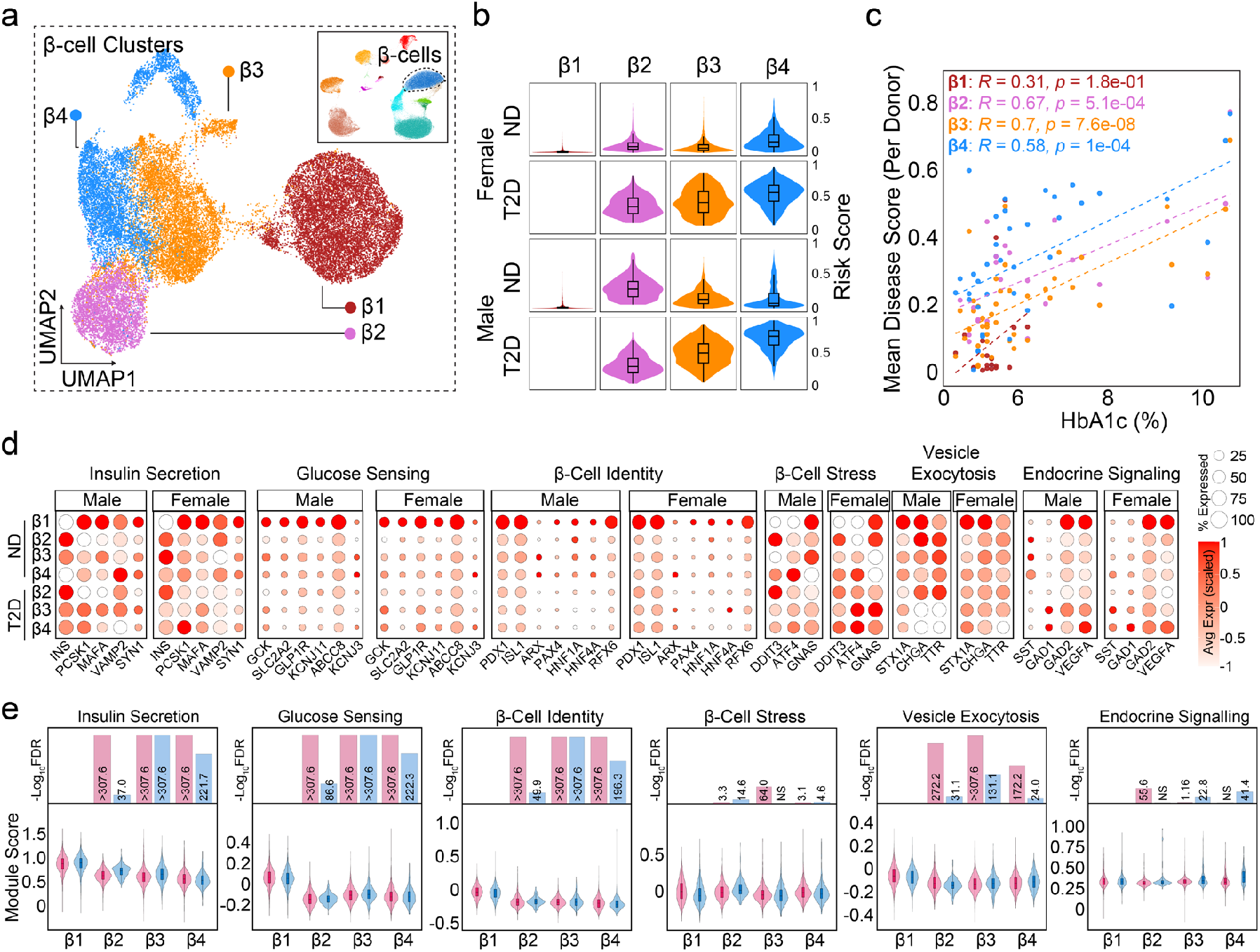
Disease scores, β cell subtype stratification, and functional gene module analysis in human islets. **a**, UMAP projection of single β cells from all donors (n=141,739 cells), colored by β cell cluster (β1-β4). Inset: UMAP highlighting β cells within the complete islet cell atlas. **b**, Violin and box plots of predicted disease risk scores for each β cell cluster (β1-β4), separated by sex and disease status (ND, T2D), illustrating subtype and sex-dependent patterns in disease risk. **c**, Scatter plots showing correlations between mean β cell disease risk scores per donor and HbA1c (%) for each β cell subtype (β1-4). Pearson correlation coefficients (R) and p-values are reported for each cluster, indicating subtype-specific associations between predicted risk and clinical glycemia. **d**, Dot plots displaying gene expression for curated functional gene sets (insulin secretion and processing, glucose sensing and metabolism, β cell development and differentiation, β cell stress and survival, vesicle trafficking and exocytosis, endocrine signaling and communication) for β cell subtypes, sex, and disease status. Dot size encodes percentage expressing cells, and color represents normalized expression. **e**, Bar and violin plots summarizing the distribution of gene module scores by β cell subtype, sex, and disease status. Numbers above bars indicate - log_10_(FDR). Statistical significance was assessed by appropriate parametric or non-parametric tests as indicated in Methods.

### Mitophagy and oxidative phosphorylation emerge as the dominant β cell programs linked to T2D

We performed tri-linear comparative pathway analysis throughout sex, disease and β cell clusters, comparing 1) ND β2-4 to ND β1, 2) T2D β2-4 to ND β1 and, 3) T2D β2-4 to ND β2-4 **(Fig. 4a)**. In pathway over representation analysis (ORA) ND β2-4 diverged from ND β1 by decreased genes involved in oxidative phosphorylation (OXPHOS), unfolded protein response (UPR), insulin secretion, glucose metabolism and lipid oxidation, and increased genes involved in stress response and cell death, independent of sex **(Fig. 4b)**. Together, results described in Fig. 3 and 4 demonstrate that the ND β1 cluster is transcriptionally superior to other β cell clusters in terms of metabolic function.

**Figure 4.**
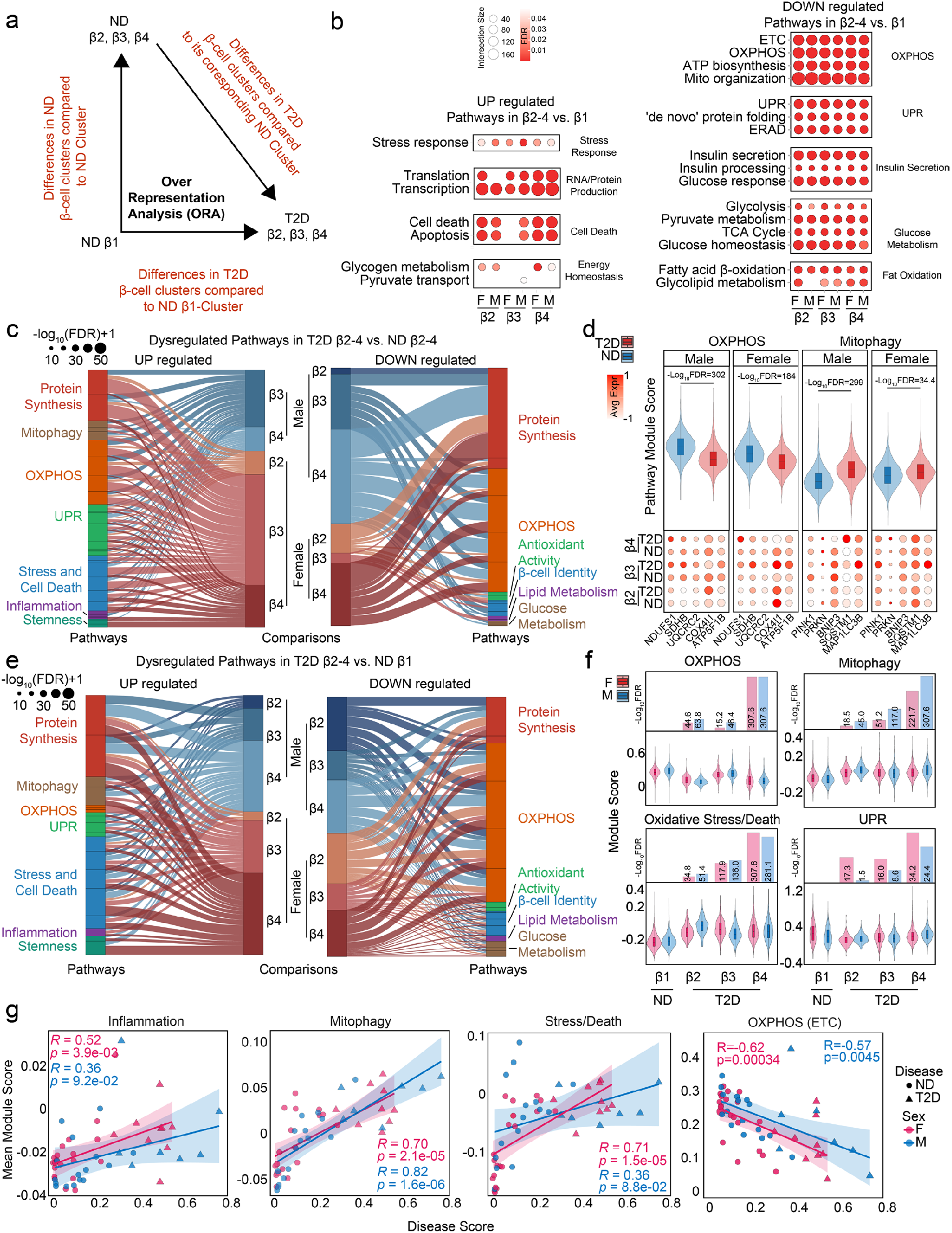
Comparative pathway analysis reveals dysregulated cellular programs in T2D β cell subtypes. **a**, Schematic overview of pathway over-representation analysis (ORA) comparing 1) ND β2-4 to ND β1, 2) T2D β2-4 to ND β1 and, 3) T2D β2-4 to ND β2-4 clusters. Arrows indicate the direction of comparison for identifying differentially regulated biological pathways between disease states and β cell subtypes. **b**, Dot plots summarizing representative pathways found to be significantly upregulated (left) or downregulated (right) in β2-4 versus β1 clusters (top panels) and in T2D versus ND β2-4 clusters (bottom panels). Dot size reflects -log10(FDR), with statistical significance determined by FDR-adjusted p < 0.05. **c**, Sankey diagrams depicting the distribution of dysregulated pathways (UP, left; DOWN, right) in T2D β2-4 clusters compared to ND β2-4 clusters male (top) and female (bottom). Pathway groups are color-coded as indicated; width of each stream reflects the strength of statistical significance (-log_10_[FDR] + 1). **d**, Violin and dot plots showing pathway module scores and select pathway specific gene expression of 5 genes, for unfolded oxidative phosphorylation (OXPHOS) and mitophagy gene sets for β-cell clusters by disease and sex. **e**, Sankey diagrams depicting the distribution of dysregulated pathways (UP, left; DOWN, right) in T2D β2-4 clusters compared to ND β1 clusters male (top) and female (bottom). Pathway groups are color-coded as indicated; width of each stream reflects the strength of statistical significance (–log_10_[FDR] + 1). **f**, Violin and dot plots showing pathway module scores for inflammation, mitophagy, oxidative stress/death and UPR for β cell clusters by disease and sex. Bars represent -log10(FDR) comparing each cluster to its β1 counterpart. **g**, Regression analysis for inflammation, mitophagy, oxidative stress/death and UPR within β cell clusters by disease and sex comparing module score and mean disease score. Details of ORA methodology and statistical testing are provided in Methods.

To explore the pathogenesis of β cell failure in T2D, we used a data visualization technique (Sankey flow diagrams) that emphasizes change in T2D β2-4 clusters from ND to T2D in which the width of the arrows is proportional to the weight of the dysregulated pathway. Compared to ND β2-4 or ND β1, T2D β2-4 showed increased stress, inflammation, UPR and mitophagy pathways and decreased protein synthesis, OXPHOS, antioxidant activity, insulin signaling, lipid metabolism and glucose metabolism pathways **(Fig. 4c, 4e** and **Supplementary Fig. 4a)**, with pathway mean module scores confirming these observations **(Fig. 4d,f** and **Supplementary Fig. 4b)**. To examine the overall impact of selected pathways **(Fig4. c-f)**, we correlated mean module scores and SnakeClassifier disease scores for all donors. For each individual donor, mitophagy showed the strongest positive correlation with disease score independent of sex (M: R = 0.7, p ≤ 2.1×10^−5^, F: R = 0.82, p ≤ 1.6×10^−6^). Only stress/death approached this in females (M: R = 0.36, p ≤ 8.8×10^−2^, F: R = 0.71, p = 1.5×10^−5^), while inflammation showed weak and UPR showed no correlation **(Fig. 4g** and **Supplementary Fig. 4c)**. OXPHOS (M: R = -0.57, p ≤ 4.5×10^−3^, F: R = - 0.62, p = 3.4×10^−4^) **(Fig. 4g)**, antioxidant activity (M: R = -0.57, p ≤ 4.5×10^−3^, F: R = -0.62, p = 3.4×10^−4^) and glucose metabolism showed negative correlation with disease score (M: R = -0.58, p ≤ 3.6×10^−3^, F: R = -0.58, p = 9.5×10^−4^) **(Supplementary Fig. 4c)**. Together, these results suggest that increased mitophagy and decreased OXPHOS are central and proportional to β cell failure in T2D.

### BlackSwanClassifier identifies mitophagy regulators *BNIP3, PINK1*, and *FUNDC1* as drivers of β cell failure and T2D severity

To gain insight into the role of mitophagy in T2D β clusters, we trained a pathway-specific classifier (BlackSwanClassifier) to identify top mitophagy candidates by mean module score. We detected mitophagy activity with AUROC = 0.72 within β clusters **(Fig. 5a)**. PTEN induced kinase 1 (*PINK1*) is a mitochondrial damage sensor involved in ubiquitin-mediated mitophagy. PINK1 accumulates on depolarized and damaged mitochondria, phosphorylates ubiquitin and recruits Parkin (*PRKN*) which then initiates ubiquitylation of mitochondrial proteins ^27,28^. In contrast, BCL2 interacting protein 3 (*BNIP3*) and FUN14 domain containing 1 (*FUNDC1*) are involved in receptor-mediated mitophagy, recruiting *MAP1LC3B*, to eliminate damaged mitochondria independently of ubiquitylation^29^. Among the top 30 ranked mitophagy candidate genes in all β clusters (by mean module score), rank aggregation of mitophagy pathway genes nominated *PINK1, BNIP3*, and *FUNDC1* as top mitophagy regulators whose dysregulation drives β cell failure associated with T2D severity **(Fig. 5b)**. *BNIP3, FUNDC1, PINK1* were downregulated while the downstream effectors of *PINK1, PRKN, SQSTM1*, and *TBK1* were upregulated **(Fig. 5c)**.

**Figure 5.**
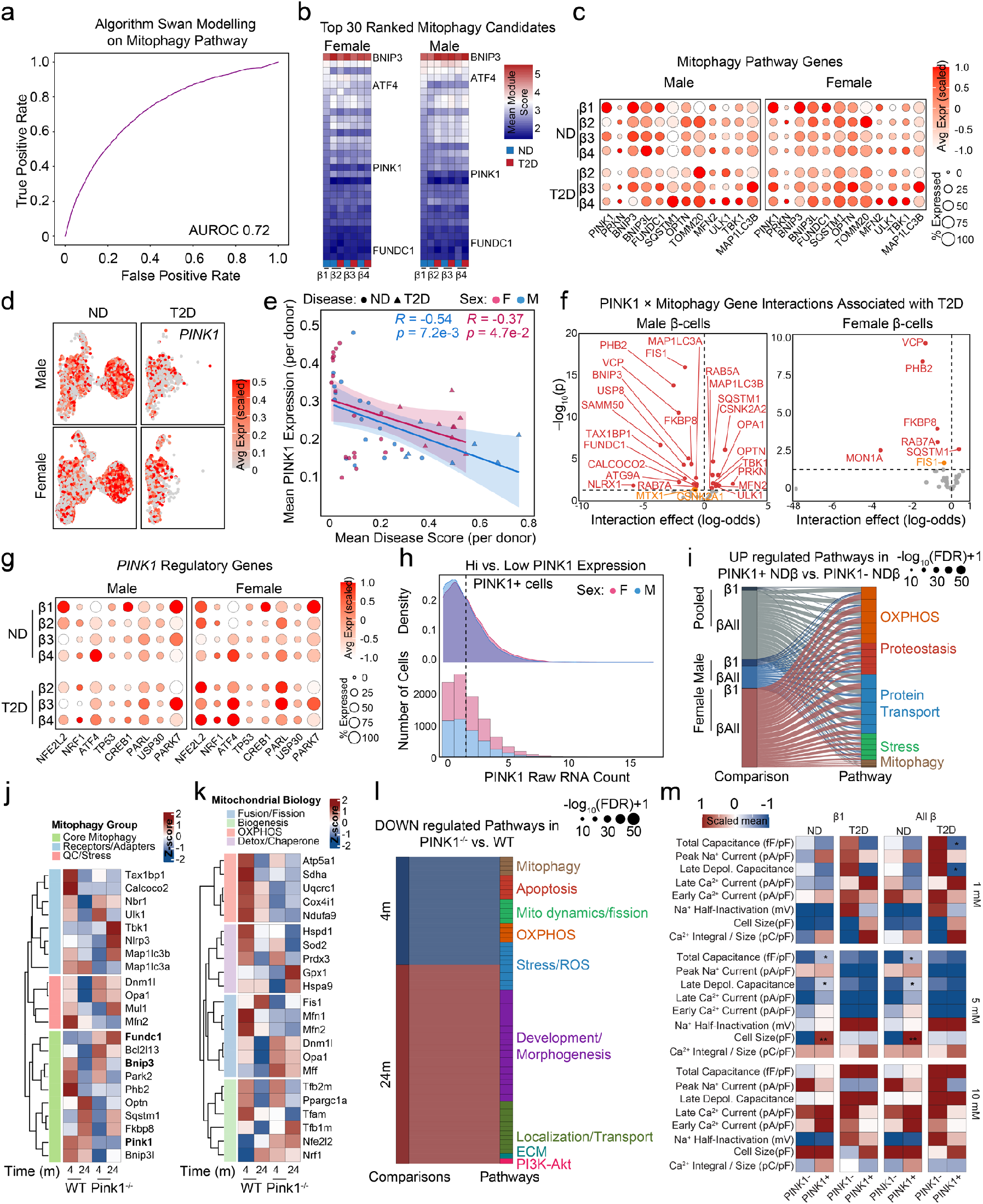
Single-cell modeling and analysis of mitophagy pathway dysregulation in T2D β cell subtypes identifies *PINK1*/*BNIP3*/*FUNDC1* as key mitophagy regulators. **a**, Receiver operating characteristic (ROC) curve illustrating performance of the Algorithm Swan logic-based model trained on mitophagy pathway features, with area under the curve (AUROC) as indicated. **b**, Heatmaps showing the top 30 ranked mitophagy candidate genes (by mean module score) in female and male β cell subtypes (ND and T2D), based on pathway module scoring. **c**, Dot plots depicting scaled average expression (color) and percent of cells expressing (dot size) key mitophagy pathway genes for β cell subtypes, stratified by sex and disease status. **d**, UMAP plots showing single-cell expression of *PINK1* for male and female β cells in ND (left) and T2D (right) groups. **e**, Correlation between mean *PINK1* expression per donor and mean disease score, colored by sex and disease, with Pearson correlation coefficients and p-values indicated. **f**, Interaction analysis linking *PINK1* with mitophagy genes identifies T2D-associated modifiers in male and female β cells (log-odds effects shown). Dashed line of y-axis marks - log10FDR of 1.301 (∼FDR=0.05). Genes in red (FDR<0.05), yellow (p<0.05). **g**, Dot plots showing average expression (color) and percentage (size) of candidate *PINK1* regulatory genes in β cell subtypes, stratified by sex and disease status. **h**, Distribution of *PINK1*^+^ cells based on raw RNA counts in pseudo-perturbation analysis, stratified by sex. **i**, Sankey diagram showing upregulated pathways (with FDR-adjusted p-values) in *PINK1*^+^ ND β cells versus *PINK1*^+^ T2D β cells, colored by pathway category as indicated. **j**, Heatmap of curated mitophagy machinery and receptor/adapter genes (grouped by function) showing scaled mean expression for WT and *Pink1*^-/-^ kidney at 4 and 24 months; columns are ordered to emphasize blocks within core mitophagy, receptors/adapters, and QC/stress modules. **k**, Heatmap for fusion/fission, biogenesis, OXPHOS, and detox/chaperone programs, highlighting coordinated remodeling of mitochondrial dynamics and respiratory complexes in Pink1^-/-^ relative to WT over time. **l**, Down-regulated pathway landscape in *Pink1*^-/-^ versus WT at 4 and 24 months; dot size encodes −log_10_(FDR)+1 and colors denote pathway families (e.g., mitophagy, apoptosis, mitochondrial dynamics/fission, OXPHOS, stress/ROS, development/morphogenesis, localization/transport, ECM, PI3K-Akt), indicating broad suppression of mitochondrial maintenance and oxidative metabolism. **m**, Patch-seq electrophysiology summary for β1 cells and all β cells (ND and T2D), stratified by *PINK1* status (*PINK1*^−^, *PINK1*^+^). Scaled means are shown for capacitance, Na^+^ and Ca^2+^ currents (peak/late), Na^+^ half-inactivation, cell size, and Ca^2+^ integral/size for stimulus steps; asterisks mark significant differences (^*^p<0.05). Patterns are consistent with altered excitability and calcium handling in *PINK1*-defined states. All p-values are two-sided; multiple comparisons were controlled with Benjamini-Hochberg FDR as detailed in methods. All p-values and FDR-adjusted values are shown as stated in each panel; statistical methods are detailed in methods.

We focused on *PINK1* as the sentinel of mitochondrial quality in β cells. At single-cell resolution, *PINK1* was abundant in ND β1 compared to other β cell clusters **(Fig. 5d)**. In individual donors, *PINK1* expression declined with mean disease score (male: R = -0.54, p = 7.2×10^−3^; female: R = -0.37, p = 4.7×10^−2^) **(Fig. 5e)**. By assessing both the magnitude of the interaction effect (log-odds) and the statistical significance of this effect (-log_10_p), we found that a larger number of *PINK1*-related mitophagy genes showed significant associations with T2D in male than female β cells **(Fig. 5f)**. These include *PHB2, VCP, FKBP8, MAP1LC3B, OPTN*, and *SQSTM1*, involved in coordinated tagging and autophagosome engagement of damaged mitochondria. In line with higher *PINK1* expression in ND β1 cells, its upstream regulators (*CREB1, NFE2L2, PARK7*) were upregulated **(Fig. 5g)**. To examine *PINK1* potential role in β cell protection, we analyzed the transcriptomes of ND β clusters with high *PINK1* (*PINK1*^+^) vs low *PINK1* expression (*PINK1*^−^) **(Fig. 5h)**. ORA pathway comparisons showed enrichment for mitophagy, OXPHOS, proteostasis and protein transport in *PINK1*^+^ compared to *PINK1*^−^ ND β clusters **(Fig. 5i)**. We profiled a *Pink1*^−/−^ mouse kidney bulk RNAseq dataset to analyze the consequences of *PINK1* loss on mitochondrial biology^30^. Heatmaps grouped by mitophagy group showed broad downregulation of core mitophagy genes **(Fig. 5j)**, and a coordinated decrease in mitochondrial OXPHOS, fusion/fission and biogenesis **(Fig. 5k)**. In addition, *Pink1*^−/−^ cells exhibited increased genes for mitochondrial UPR activation alongside increased inflammation, DNA damage, oxidative-stress and apoptosis **(Supplementary Fig. 5a-b)**. Over-representation analysis of downregulated pathways confirmed these effects, highlighting mitophagy, apoptosis, mitochondrial dynamics/fission, OXPHOS, and stress/ROS as the most affected categories, with additional suppression of developmental/ECM and PI3K-AKT programs **(Fig. 5l)**.

We next leveraged human islet Patchseq^25,31^ (combinatorial electrophysiological measurements and scRNAseq), confirming increased exocytotic capacity of ND *PINK1*^*+*^ β cells (as total capacitance) at 5 mM glucose, which results mostly from an increased late exocytotic response thought to reflect insulin granule recruitment **(Fig. 5m)**. We also observed similar results at 10mM glucose in β cells having high MQI **(Supplementary Fig.5c-d)**. There is some indication, particularly in the high MQI ND β cells **(Supplementary Fig.5d)**, that this increased exocytosis may be driven by greater Ca^2+^ entry and shifts in excitability due to altered Na^+^ channel activity. These responses are largely lost in the T2D β cells.

### Androgen action shifts β cell metabolism toward mitophagy

To gain insight on the cause of the greater mitophagy gene correlation with T2D in males (**Fig. 5f)**, we mapped male human β cells treated with dihydrotestosterone (DHT)^32^ onto the β cluster map (β1-β4; **Fig. 6a**). DHT produced a broad induction of *BNIP3* **(Fig. 6b)**. We used Seurat’s module score function to model mitochondrial function. Bioenergetic module showed a DHT-induced shift from OXPHOS to glycolysis (reduced OXPHOS/glycolysis and OXPHOS/TCA; **Fig. 6c**), consistent with our previous report^32^. For all β cell clusters DHT stimulation increased mitophagy (module score, p = 0.0012; **Fig. 6d**), without difference between clusters **(Fig. 6e)**. Thus, *AR* activation in β cells increases mitophagy. To investigate if mitophagy influences inflammation we evaluated the expression of proinflammatory genes and saw an increase in T2D in both sexes other than *IRF3/7, HMGB1* and *IKBKG* which were elevated in ND β1 **(Supplemental Fig. 6a)**. DHT decreased proinflammatory gene expression vs controls **(Supplemental Fig. 6b)**.

**Figure 6.**
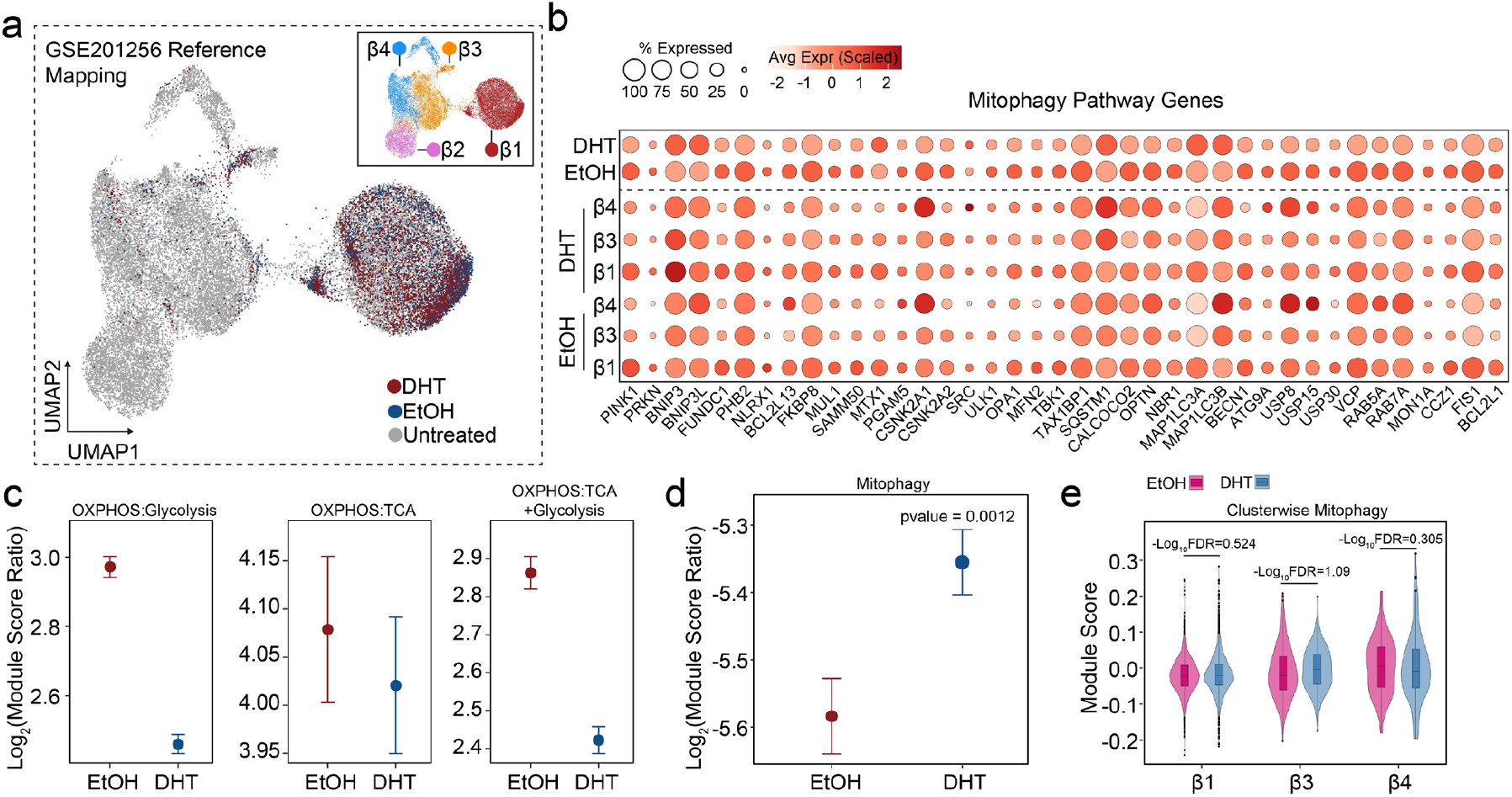
DHT reprograms β cell energetics and mitophagy. **a**, Reference mapping of our dataset to the human islet DHT dataset GSE201256. UMAP shows mapped cells colored by condition (DHT, EtOH, Untreated). Inset: reference β cell subclusters (β1-4) used for cluster labels. **b**, Dot plot of curated mitophagy genes for β cell subclusters under DHT and EtOH. Dot size indicates the percentage of cells expressing the gene; color denotes scaled average expression. Dashed line separates conditions. **c**, Bioenergetic balance quantified as log_2_ module-score ratios: OXPHOS:Glycolysis (left), OXPHOS:TCA (middle), and OXPHOS:(TCA+Glycolysis) (right). Points indicate group means; error bars show 95% CIs. Relative to EtOH, DHT shifts β cells toward lower OXPHOS scores versus glycolytic/TCA programs. **d**, Global mitophagy module scores by condition (EtOH vs DHT); points are means with 95% CIs. P value from a two-sided Wilcoxon test (p = 0.0012). **e**, Cluster-wise distribution of mitophagy scores for β1, β3, and β4 under EtOH (magenta) and DHT (blue). Violins show per-cell distributions with embedded boxplots (median and IQR). Numbers above brackets report - log_10_(FDR) from Benjamini-Hochberg corrected two-sided tests. Module scores were computed from pre-specified gene sets (see Supplementary Methods); ratios were calculated on per-cell scores before aggregation by condition/cluster. EtOH denotes vehicle control; DHT denotes dihydrotestosterone.

### Mitochondrial function index links mitophagy/proteostasis to β cell metabolic fitness

Kolmogorov Arnold neural networks (KANN) approximate arbitrary multivariate maps via additive learned univariate functions, making them well suited to the analysis of scRNA-seq data, where complex geometric interactions within multiple pathways can be modeled into a singular interpretable codex.^33-37^ We reasoned that transcriptional networks driving mitochondrial function can be modelled *in silico* using a KANN. We modeled mitochondrial UPR, OXPHOS, mitophagy, and biogenesis core pathways into a single mitochondrial function index (MFI). For comparative purposes, we also generated a mitochondrial quality index (MQI), which is the mean of pathway module scores used to train the KANN. The post training estimated MFIs from single-cell transcriptomes showed a strong correlation with module-driven observed MQI (R^2^ = 0.934; RMSE = 0.436) (**Fig. 7a,b)**. Permutation-importance profiles were highly robust across random seeds (Spearman ρ = 0.999-1) with strong concordance in the top 30 ranked features (Jaccard 0.935-1) **(Supplemental Fig. 7a)**. MFI feature robustly weighted mitophagy machinery and mtUPR/proteostasis, with *PINK1*/*PRKN*/*SQSTM1* axis components and *BNIP3* among the strongest contributors, followed by fusion/fission and antioxidant modules (**Fig. 7c)**. Overall, MFI accurately mapped higher scoring (MFI>1) for ND vs poorly scoring (MFI<1) T2D cells **(Fig. 7d)**. Specifically, higher MFI (>1) were observed in ND β1 compared to ND β2-3 and T2D β2-4 in both sexes, except T2D β3 in females (**Fig. 7e-f)**. Individual donors with low MFI trended with increasing HbA1c levels (**Fig. 7g**). T2D β3 with high MFI (>1) exhibited increased OXPHOS, UPR and mitophagy pathway activation compared to T2D β3 with low MFI (<1) **(Supplementary Fig 7b)**. Differential expression within untreated β1 revealed that high-MFI cells upregulated OXPHOS/ETC, ATP biosynthesis and mtUPR (FDR<0.05) and downregulated stress/injury, secretory and cell death pathways (**Fig. 7h-j)**.

**Figure 7.**
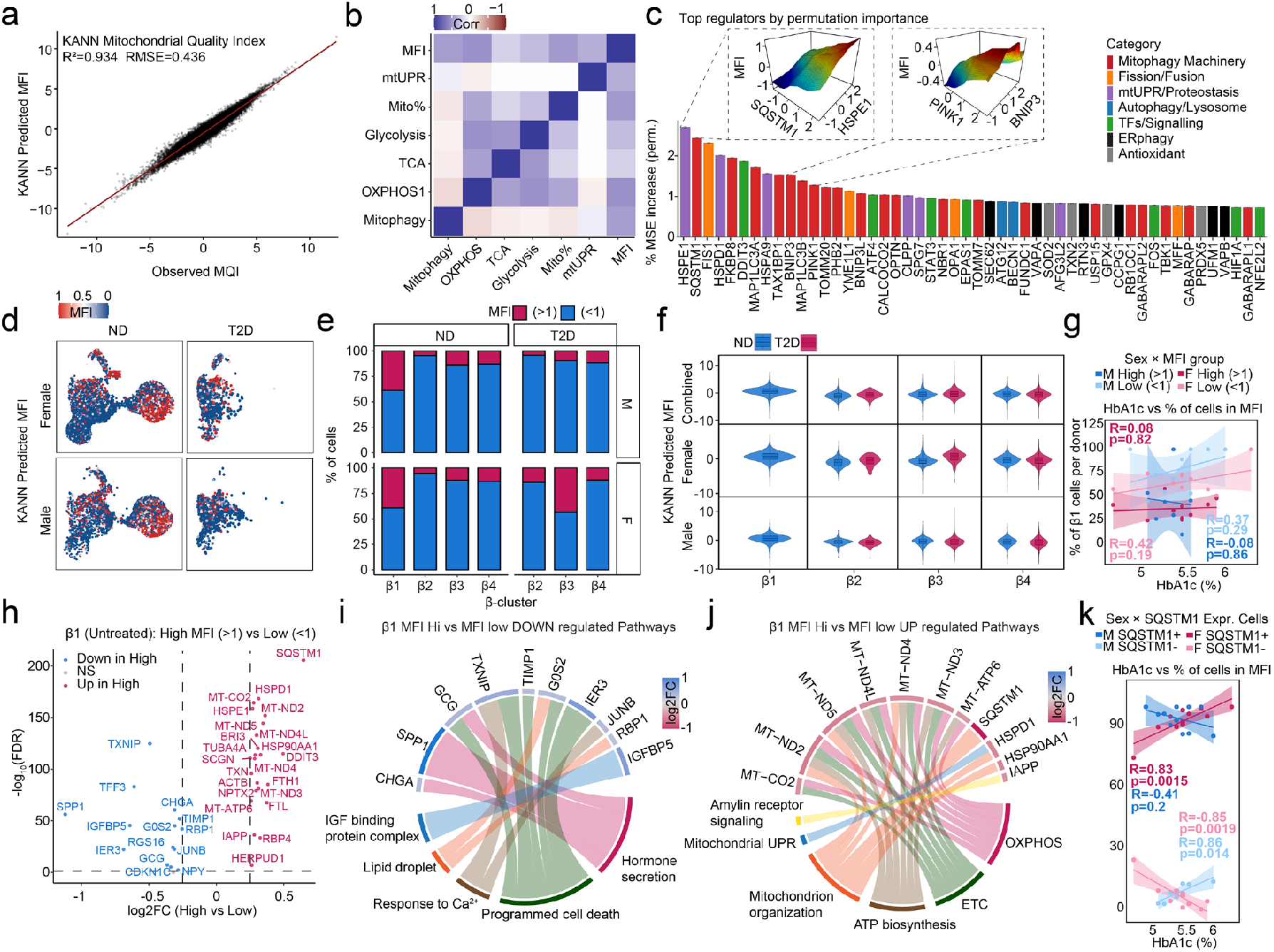
Interpretable KANN-derived mitochondrial fitness (MQI/MFI) captures β cell heterogeneity, regulators, and clinical links. **a**, Performance of the Kolmogorov-Arnold Neural Network (KANN) model predicting the mitochondrial fitness index (MFI) from single-cell gene expression. Scatter shows observed MQI versus KANN-predicted score; line is identity (R^2^ = 0.934; RMSE = 0.436). **b**, Correlation structure between the KANN score (MFI/MQI) and pathway module scores (mitophagy, OXPHOS, OXPHOS:TCA ratio, glycolysis, TCA, mitochondrial content, mtUPR). **c**, Permutation importance of candidate regulators. Bars indicate % increase in prediction MSE when a regulator is permuted; colors group functional classes (mitophagy machinery, fission/fusion, mtUPR/proteostasis, autophagy/lysosome, TFs/signaling, ER-phagy, antioxidant). Insets show pairwise partial-dependence surfaces for exemplar genes. **d**, UMAP view of β-cells colored by KANN-predicted MFI for non-diabetic (ND) and T2D donors, split by sex. **e**, Fraction of cells per β-cell subcluster (β1-4) with high versus low MFI in ND and T2D, stratified by sex. Thresholds: High (>1) and Low (<1). **f**, Distribution of KANN-predicted MFI by β cell subcluster for ND vs T2D (combined and stratified by sex). Violins show per-cell distributions with embedded boxplots (median and IQR). **g**, Donor-level association between HbA1c and the proportion of β1 cells in high or low MFI groups, stratified by sex. Lines show linear fits; annotations report Pearson R. **h**, Differential expression in β1 (Untreated) comparing MFI-High (>1) vs MFI-Low (<1). Volcano plot highlights representative down (blue) and upregulated (red) genes. **i**, Enriched pathways among genes down-regulated in β1 MFI-High vs MFI-Low. Chord diagram links genes to pathway terms; segments are colored by log_2_ fold-change. **j**, Enriched pathways among genes up-regulated in β1 MFI-High vs MFI-Low, emphasizing OXPHOS/ETC, ATP biosynthesis, mitochondrial organization, mtUPR, and amylin-receptor signaling. **k**, HbA1c versus fraction of β1 cells in high-MFI per donor, stratified by sex and *SQSTM1* expression status (*SQSTM1*^+^ /*SQSTM1*^-^). Lines denote linear fits; Pearson R and two-sided P values are shown.

The second hit in our KANN classification, *SQSTM1* or p62 is a critical mitochondrial capping protein which is recognized by the autophagosome via *MAP1LC3B*. Increasing HbA1c correlated with higher percentage of *SQSTM1*^+^ β cells in females (R=0.83, p=0.0015), while conversely higher HbA1c correlated to higher percentage of *SQSTM1*^−^ cells in males (R=0.86, p=0.014, **Fig. 7k**). Thus, the MFI framework demonstrates a mitophagy proteostasis axis, driving β cell metabolic fitness which decreases in T2D in both sexes **(Fig. 8)**.

**Figure 8.**
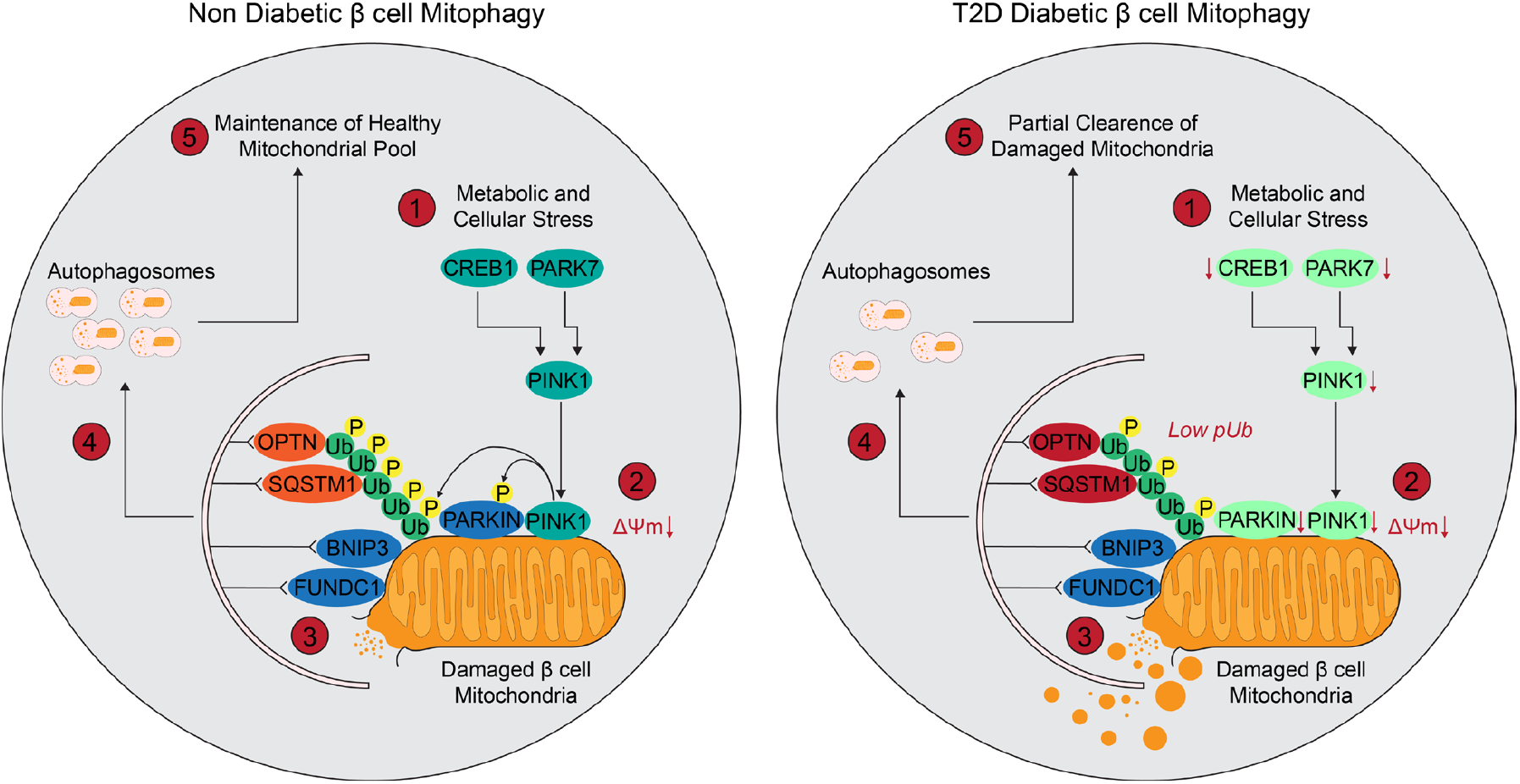
Model of impaired mitophagy in β cells during type 2 diabetes development. **(1)** In healthy β cells, metabolic and cellular stress induce CREB1 and PARK7, which transcriptionally activate PINK1. In T2D β cells, both CREB1 and PARK7 are downregulated, reducing PINK1 induction. **(2)** Upon mitochondrial depolarization (ΔΨm↓), stabilized PINK1 phosphorylates both PARKIN (Ser65) and ubiquitin (Ub; Ser65), generating phospho-ubiquitin (p-Ub) chains that amplify PARKIN activation. In T2D, reduced PINK1 activity leads to fewer p-Ub-modified chains and diminished PARKIN activation. **(3)** Activated PARKIN ubiquitinates outer mitochondrial membrane substrates, which are recognized by mitophagy receptors. In non-diabetic cells, OPTN, SQSTM1, BNIP3, and FUNDC1 coordinate to recruit the phagophore membrane. In T2D, OPTN and SQSTM1 expression increase, but act mainly on non-phosphorylated Ub, reflecting a compensatory receptor-driven mitophagy response. **(4)** In non-diabetic β cells, these receptor complexes enable efficient autophagosome formation and clearance of damaged mitochondria. In T2D, this process is incomplete, with fewer mitochondria successfully engulfed. **(5)** Consequently, healthy β cells maintain a renewed mitochondrial pool, whereas diabetic β cells show partial clearance and accumulation of damaged mitochondria, impairing mitochondrial quality control and β-cell function.

## Discussion

The methodological innovation of this study lies in combining rule-based logic models with pathway-aware constraints and Kolmogorov-Arnold neural networks (KANN) to produce interpretable classifiers of β cell states. Existing approaches for single-cell disease modeling including random forests, gradient boosting, and deep neural networks provide high accuracy but limited transparency, often operating as black boxes with little mechanistic interpretability^21,22^. Recent innovations in graph neural networks and transformers have improved annotation and clustering^19,20^, yet these methods remain largely correlative and do not resolve causal gene programs. Our framework advances beyond these approaches by (i) enforcing sparse Boolean rules (SnakeClassifier) that yield human-interpretable thresholds, (ii) constraining models to biological pathways (BlackSwanClassifier) to preserve mechanistic plausibility, and (iii) leveraging KANN to collapse high-dimensional pathway activity into a unified mitochondrial fitness index, overcoming the problem of dimensionality while retaining interpretability^33,36^. Collectively, these advances produce accurate disease state prediction while simultaneously exposing the molecular pathogenesis of β cell failure.

Using this modeling framework, we identified mitophagy as a central determinant of β cell failure in T2D. In both sexes β cells exhibit suppressed mitochondrial respiration and a decline in *PINK1* expression, suggesting failure of ubiquitin-mediated mitophagy. In addition, male β cells displayed androgen-induced upregulation of *BNIP3* and metabolic reprogramming toward glycolysis. Thus, both sexes converged on a shared state characterized by elevated mitophagy and reduced oxidative phosphorylation, tightly correlated with disease burden.

*PINK1* is a sentinel of mitochondrial quality control that shields β cells from oxidative collapse. Decline in PINK1 coupled with sustained activity of *BNIP3* and *FUNDC1* results in unchecked mitochondrial clearance, reduced respiratory capacity, dismantling of the fission/fusion/biogenesis axis, and a shift toward proteotoxic stress, secretory failure, and apoptosis. These observations converge with prior reports that mitophagy protects β cells from inflammatory and metabolic stress^29,38,39^ and that *BNIP3* mediated mitophagy can protect human β cells under diabetogenic conditions^40^. Together with studies implicating *PINK1-PRKN* signaling in mitochondrial surveillance^27,28^, our results suggest that defective coordination between sensors and degradative effectors of damaged mitochondria is a core vulnerability in β cell failure.

Looking forward, this work illustrates how mechanistically grounded machine learning can generate disease models that both stratify risk and expose candidate regulators. While our current framework was trained on transcriptomic features, its extension to multiomic modalities (chromatin accessibility, proteomics, metabolomics) will be critical in refining disease trajectories^13,14^. Moreover, functional validation of computationally nominated regulators across genetic backgrounds and *in vivo* systems remains essential. More broadly, our work illustrates how interpretable predictive single cell ML models can transform precision medicine by tracing T2D risk back to specific molecular programs. Utilizing this approach allowed us to determine not only which β cells are vulnerable, but also why, and to uncover gene networks that may be targeted to restore cellular resilience.

## Materials and Methods

### Mathematical Foundations of SnakeClassifier and BlackSwanClassifier

The SnakeClassifier and BlackSwanClassifier derive their theoretical basis from the use of NP-complete structures specifically, the Boolean Satisfiability Problem (SAT) to generate machine learning insights for classification tasks. At the core of Algorithme.ai’s research lies a mathematical proposition pertaining to Supervised Machine Learning.

Let X denote a truth vector of size m and let A be a binary (0-1) matrix of appropriate dimensions such that no two rows of A are identical. Under this premise, there exists a SAT instance that can be constructed in *O(nm*^*2*^) time and that achieves a perfect fit on the training dataset.

### The Dana Theorem: Constructive SAT Representation of Supervised Boolean Functions

Let *A* ∈ {0,1}^*n*×*m*^ be a Boolean matrix with distinct rows and let *X* ∈ {0,1}^*n*^ be a Boolean label vector. Define the sets of positive and negative indices:

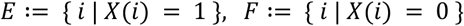

Then there exists a CNF formula *φ* with at most |*F*| ≤ *n* clauses and |*E*| · |*F*| ≤ ⌊*n*^2^/4⌋ literals, constructible in time *O*(*m* · |*E*| · |*F*|) ⊆ *O*(*mn*^2^), such that:

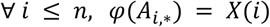

#### Proof

For any pair (*i, j*) ∈ *F* × *E*, define a conflict index:

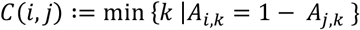

Because all rows of A are distinct, such an index always exists, and computing it costs *O*(*m*). The total number of pairs (*i, j*) is |*F*| · |*E*|.

#### Step 1. Constructing discriminative literals

For each pair (*i, j*) ∈ *F* × *E*, define a separating literal:

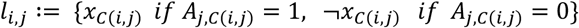

By construction, *l*_*i,j*_(*A*_*j*,∗_) = 1 and *l*_*i,j*_(*A*_*i*,∗_) = 0.

#### Step 2. Clauses for the negative population

For each negative index *i* ∈ *F*, define the discriminative clause: *C*_*i*_ := ⋁_*j*∈*E*_ *l*_*i,j*_

The cost of constructing one clause is *O*(*m* · |*E*|), so building all of them is *O*(*m* · |*E*| · |*F*|).

#### Step 3. Global conjunction

Define the full CNF formula as: *φ* := ⋀_*i*∈*F*_ *C*_*i*_ Thus, the number of clauses is |*F*| ≤ *n*, and the total number of literals is |*E*| · |*F*| ≤ ⌊*n*^2^/4⌋.

#### Step 4. Verification of correctness

For any positive example *j* ∈ *E*: For every *i* ∈ *F, l*_*i,j*_(*A*_*j*,∗_) = 1; hence each *C*_*i*_(*A*_*j*,∗_) = 1. Consequently, *φ*(*A*_*j*,∗_) = 1 = *X*(*j*). For any negative example *i* ∈ *F*: For all *j* ∈ *E, l*_*i,j*_*A*_*i*,∗_ = 0, so *C*_*i*_(*A*_*i*,∗_) = 0. Therefore, *φ*(*A*_*i*,∗_) ≤ *C*_*i*_(*A*_*i*,∗_) = 0 = *X*(*i*).

#### Step 5. Complexity

Each conflict *C*(*i, j*)can be found in *O*(*m*) time, and there are |*E*| · |*F*| such pairs. Hence the total construction time is *O*(*m* · |*E*| · |*F*|) ⊆ *O*(*mn*^2^). Edge cases

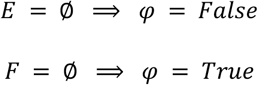

#### Corollary 1 (Dual DNF form)

The dual construction swapping the roles of *E* and *F* produces a DNF formula ψ that satisfies the same property:

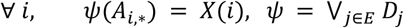

### Clause-Level Interpretability in the BlackSwanClassifier Framework

*φ* exactly reproduces the target labeling *X* on the finite dataset *A* and can be constructed in polynomial time. Hence, any finite Boolean indicator function admits an equivalent SAT instance constructible in *O*(*mn*^2^) which is precisely the claim of the Dana Theorem. In the initial formulation, SAT instances were generated separately to represent both True and False outcomes. A confidence measure was then obtained by subtracting the number of False instances from the number of True instances across N=50 samples, producing a coarse confidence percentage. However, this approach exhibited limited signal granularity.

To address this limitation, the SnakeClassifier was introduced. It constructs a table of lookalikes, each directly referencing the training basis. The algorithm begins by assuming that every datapoint is a potential lookalike of the query instance. For each activated clause in the SAT indicator corresponding to a True outcome, it removes lookalikes that do not satisfy the clause (i.e., False lookalikes). Conversely, for clauses indicating a False outcome, it removes lookalikes that satisfy the clause (i.e., True lookalikes). This process is applied iteratively across multiple layers of lookalikes. The resulting confidence score is computed as a/b, where a is the number of remaining True lookalikes and b is the total number of lookalikes considered.

The BlackSwanClassifier introduces further refinement by emphasizing the fundamental SAT construct the clause. Instead of relying on aggregate instance subtraction, it systematically constructs discriminative clauses that distinguish the True population with respect to the False elements, and vice versa. This clause-level modeling substantially improves computational efficiency and discriminative power.

Empirically, this advancement reduced model compilation time from several hours to approximately 350 seconds on a standard personal computer, using a pure Python implementation trained on a dataset comprising 5,000 samples.

## Data sources and cohort assembly

We analyzed single cell RNA sequencing (scRNAseq) profiles from 52 human pancreatic islet donors (GSE266291), including non-diabetic (ND) and type 2 diabetic (T2D) individuals (141,739 cells). Metadata included donor, sex, cell type, and clinical measures (e.g., HbA1c). For androgen-perturbation mapping, cells were transferred to the human islet atlas GSE201256 to obtain harmonized β cell subcluster labels (β1-4). All downstream analyses preserved donor, sex, and disease labels to prevent leakage.

### Preprocessing and quality control

Standard single-cell QC was applied (minimum detected genes, mitochondrial RNA percentage, and doublet screening where applicable). Counts were normalized and variance-stabilized (e.g., log-normalize or SCTransform), highly variable genes were selected, and donor effects were corrected when needed using anchor-based integration via Harmony^12^. PCA and UMAP were used for visualization only. Cell type assignments were harmonized throughout donors using canonical markers and prior annotations^3^.

### Self-supervised pretraining and classifier training

To build cell-level disease state predictors, we first performed self-supervised representation learning on the full atlas while constraining supervised labels to <3% of cells. We trained three model families on these embeddings and/or selected gene features: Gradient Boosting, Random Forest, and a logic-based interpretable model (SnakeClassifier). For the logic model, we curated 50 most variable genes showing expression in atleast 50% cells, and 3 other features including disease status, sex and cell type. We utilized 5,000 representative (T2D/ND) cells to learn sparse Boolean clauses (thresholded literals combined by AND/OR), aggregated by a calibrated logistic layer to yield per-cell disease risk probabilities, termed as risk scores. Training respected donor-grouped splits to ensure sex-aware, donor-generalizable evaluation.

### Cell-level disease scoring, thresholds, and evaluation

Each model produced a continuous disease risk score for every cell in the atlas. We visualized scores on UMAPs and re-clustered to annotate inferred disease states. Performance was assessed on held-out donors using ROC curves and AUROC; operating points were selected by scanning Youden’s index. We summarized mean AUROC by sex and used a “weighted-winner” analysis, in which each subgroup (sex × ancestry × cell type) contributed proportional evaluation weight, to compare Algorithm Snake with Gradient Boosting and Random Forest. Concordance between predicted and observed T2D fractions was computed per cell type and sex. Donor-level biological relevance was tested by correlating mean cell-level disease scores with HbA1c. Unless noted, tests were two-sided; AUROC differences used DeLong’s test; correlations report Pearson’s R with bootstrap CIs and FDR correction where multiple comparisons were performed.

### β cell extraction, subtype stratification, and clinical association

β cells were isolated from the atlas, embedded with PCA/UMAP, and clustered at low resolution into four robust subtypes (β1-4). Predicted disease risk distributions were compared within β1-4 using violin-with-box overlays and stratified by sex and ND/T2D status. For each subtype, donor-level mean β cell disease scores were correlated with HbA1c, reporting Pearson’s R and FDR-adjusted p-values to highlight subtype-specific clinical links.

### Functional module scoring and coupling to risk

Curated gene sets captured insulin secretion/processing, glucose sensing/metabolism, β cell development/differentiation, stress/survival, vesicle trafficking/exocytosis, and endocrine signaling/communication. Per cell module scores (AddModuleScore^26^) were summarized as means with 95% CIs by β cell subtype, sex, and disease status (dot/violin/bar plots). Group differences used parametric or non-parametric tests with Benjamini-Hochberg FDR control. We quantified coupling between disease risk and biology by correlating donor-level mean module scores with (i) mean β-cell disease scores and (ii) HbA1c, stratified by sex and β cell subtype (Pearson R, FDR-adjusted p-values).

### Pathway over-representation analysis

To compare disease programs for β cell subtypes, we performed differential expression for (i) β2-4 versus β1 in ND donors and (ii) T2D versus ND within β2-4, using a Wilcoxon rank-sum test with minimum expression fraction and log_2_ fold-change thresholds. Significant genes (FDR < 0.05) were submitted to ORA (e.g., g:Profiler^41^/GO Biological Process^42,43^) using an atlas-matched background. Dot plots summarize representative up- and down-regulated pathways (dot size encodes -log_10_[FDR] or intersection size, as indicated). Sankey diagrams display dysregulated program flow, with comparison to pathway edge weights set to -log_10_(FDR)+1 and pathway to gene links reflecting representative intersections; pathway families (e.g., translation, ER stress/proteostasis, lipid homeostasis, metabolism) were color-grouped to match figure annotations.

### Mitophagy focused modeling and readouts

A pathway-focused logic variant (BlackSwanClassifier) was trained on mitophagy features to quantify discriminative capacity (AUROC on held-out donors). The mitophagy panel comprised core *PINK1*-*PRKN* machinery, *TOMM* import components, adaptors (*SQSTM1*/*OPTN*/*CALCOCO2*/*TAX1BP1*), receptor mediated routes (*BNIP3*/*BNIP3L*/*FUNDC1*/*PHB2*/*FKBP8*/*BCL2L13*), and regulators (e.g., E3/DUBs, fission/fusion). Heatmaps highlighted top candidates by mean module score within β clusters, sex, and disease; dot plots showed scaled average expression and percent expressing; UMAP feature plots displayed *PINK1*. Donor-level mean *PINK1* expression was correlated with mean disease score (sex-stratified). A pseudo-perturbation analysis profiled *PINK1*-positive fractions by sex. Sankey diagrams contrast pathway enrichments between *PINK1*^+^ ND and *PINK1*^+^ ND cells with FDR adjustment, using the same edge-weighting scheme described above. Bulk RNAseq datasets of PINK1KO mice kidneys (GSE217775^30^) were used to study the effect of PINK1 loss. 4m and 24m data was stratified for pathways related to mitochondrial function and biology, and subsequent differentially expressed genes (DESeq2) were used to perform ORA.

### Androgen signaling analysis and atlas mapping

For androgen exposure, cells (EtOH vehicle, DHT: GSE201256^32^) were mapped to the GSE266291^3^ reference using transfer anchors to obtain β1-4 labels. We visualized curated mitophagy genes by condition and subcluster (dot size = % expressing; color = scaled mean). Cellular bioenergetics were summarized via per-cell module-score ratios: OXPHOS:Glycolysis, OXPHOS:TCA, and OXPHOS:(TCA+Glycolysis); group means with 95% CIs are shown. Global mitophagy module scores (EtOH vs DHT) were compared with a two-sided Wilcoxon test. Cluster-wise mitophagy score distributions used violins with embedded boxplots; significance bars report -log_10_(FDR) from Benjamini Hochberg-corrected tests. Ratios were computed on per-cell scores before aggregation; EtOH denotes vehicle control; DHT denotes dihydrotestosterone.

### Patch-Seq

Existing patch-seq data^25,31,44^ were from cells cultured in low-glucose (5.5 mmol/L) DMEM with L-glutamine, 110 mg/L sodium pyruvate, 10% FBS, and 100 U/mL penicillin/streptomycin for approximately 1-3 days prior to recordings. Whole-cell and perforated patch-clamp electrophysiology, cell collection, library preparation, sequencing and cell-typing were as described^25,31,44^. Reads were aligned to the human genome (GRCh38 with ERCC sequences) using STAR^45^, and gene counts quantified by htseq-count (intersection-nonempty) using Ensemble89 GTF annotations^46^. Experiments were performed with with 1, 5 or 10 mM glucose in the bath solution. Analyses included cells from donors with no diabetes (ND) or type 2 diabetes (T2D).

### Kolmogorov-Arnold Network for mitochondrial fitness

A Kolmogorov-Arnold Neural Network (KANN) was trained to predict a mitochondrial fitness index (MFI) from single-cell expression using spline/basis functions with L1/L2 regularization and early stopping; donor grouped splits ensured out of sample evaluation. Performance was summarized by R^2^ and RMSE on held out donors. Associations between KANN predicted MFI and pathway modules (mitophagy, OXPHOS, OXPHOS:TCA, glycolysis, TCA, mitochondrial content, mtUPR) were assessed via Pearson correlations. Regulator contributions were quantified by permutation importance (% increase in prediction MSE), and pairwise partial dependence plots were generated for exemplar regulators. UMAPs were colored by MFI, split by sex and disease. Cells were dichotomized into High (>1) and Low (<1) MFI to summarize subtype fractions (β1-4) and to compare ND vs T2D (sex-stratified). Within β1 (Untreated), differential expression contrasted MFI-High vs MFI-Low; volcano and chord/ORA diagrams summarized enriched programs (e.g., OXPHOS/ETC, ATP biosynthesis, mitochondrial organization, mtUPR, amylin receptor signaling). Donor-level HbA1c associations were also tested for the fraction of β1 cells with high MFI, additionally stratified by *SQSTM1* status.

### Kolmogorov-Arnold Representability in Neural Form

The mitochondrial fitness index (MFI) model is implemented as a Kolmogorov-Arnold Neural Network (KANN), trained to learn a continuous mapping *i*: *R*^*P*^ → *R* from a β-cell’s gene expression vector *x* = (*x*_1_, …, *x*_*P*_) to a scalar mitochondrial fitness score. By the Kolmogorov-Arnold representation theorem, any multivariate function can be expressed as a finite superposition of univariate functions and linear combinations:

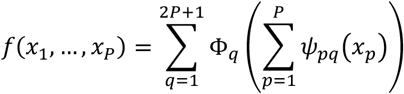

The KANN operationalizes this principle through shallow compositional layers that mix linear transformations and learned univariate spline corrections:

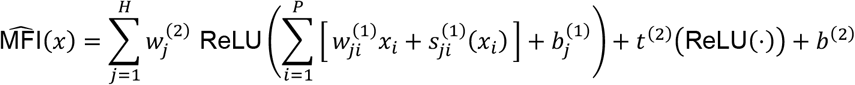

Here, *H* denotes the hidden width, *w*^(1)^, *w*^(2)^and *b*^(1)^, *b*^(2)^are trainable linear parameters, and 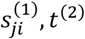 are spline-based univariate nonlinearities. This structure ensures that complex mitochondrial response surfaces can be represented as finite superpositions of interpretable, low-dimensional components.

### Spline Basis as a Stable Univariate Expansion

Each nonlinear term 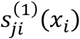 is defined as a linear combination of hat-basis (piecewise-linear) functions:

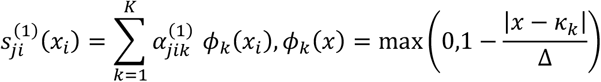

where knots *k*_*k*_ are uniformly spaced on [−3,3] and Δ = *k*_*k*+1_ − *k*_*k*_. By registering knots as fixed buffers and learning only coefficients 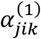, the model achieves smooth univariate responses with stable gradients, avoiding the instability of free knot optimization. In implementation, each 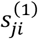 corresponds to a *KANSpline1D* module in R/torch, ensuring deterministic univariate correction functions per feature.

### Donor Grouped Evaluation and Generalization

To prevent overfitting individual cell identities, model training and validation are performed using donor grouped partitions. Let 𝒟_train_ and 𝒟_test_ denote disjoint donor sets. By evaluating model performance (R^2^, RMSE) exclusively on unseen donors, the KANN-MFI framework estimates between donor generalization, ensuring that the learned function *i*_Θ_ captures invariant biological structure rather than donor specific variance. This design grounds the MFI as a donor agnostic quantitative phenotype of mitochondrial resilience.

### Permutation Importance as a Measure of Predictive Necessity

Feature relevance is quantified through permutation importance. For a fitted model *i*_Θ_, let 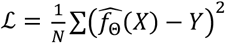 denote baseline mean-squared error, and 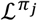 the error after independently permuting column *j*. The importance of feature *j* is defined as 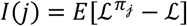 which is non-negative and equals zero if and only if the feature is conditionally irrelevant under the joint distribution of *X, Y*. This formulation aligns interpretability with predictive necessity: the higher the increase in MSE, the more indispensable the gene for accurate MFI estimation. In practice, multi-seed and multi-repeat permutation runs yield mean ± SD consensus importance scores across all features.

### Hierarchical Interpretability in the MFI Framework

Interpretability arises intrinsically from the KANN’s additive, univariate structure. At the local level, each spline function 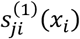 describes how the expression of a specific gene modulates the hidden response driving mitochondrial fitness directly visualized through spline gallery plots. At the global level, permutation-based importance identifies which regulators (e.g. *PINK1, BNIP3, YME1L1*) most consistently influence predicted MFI across cells and donors. Together, these layers yield a mechanistic representation of mitochondrial function, one that is not inferred *post hoc* but encoded within the network’s mathematical structure. By combining fixed knot spline expansions, donor aware validation, and necessity-based feature ranking, the MFI acts as both a predictive and mechanistic model quantitatively linking gene level expression patterns to emergent mitochondrial performance.

### Statistics, multiple testing, and visualization

Unless specified, tests were two-sided. AUROC differences used DeLong’s test; threshold selection used Youden’s index. For module and pathway analyses, Benjamini–Hochberg FDR controlled multiple hypotheses. Correlations reported Pearson’s R with FDR-adjusted p-values for families of tests (e.g., β clusters, sex, or pathways). UMAPs, ROC curves, threshold scans, violin/box plots, dot plots, bar charts, heatmaps, partial-dependence surfaces, Sankey, and chord diagrams were generated with standard plotting libraries. For Sankey, comparison to pathway edge widths reflect -log_10_(FDR)+1; for dot plots, dot size encodes either intersection size or % expressing and color denotes scaled mean or -log_10_(FDR), as stated on panels.

### Reproducibility and environment

Random seeds were fixed, and donor-grouped splits were used throughout to prevent leakage. Intermediate objects (risk scores, β cluster labels, differential expression tables, ORA outputs, module scores, KANN predictions) were saved to structured directories to enable deterministic figure re-rendering without recomputation.

## Data Availability

Single-cell RNA-sequencing data of T2D and ND islets is available on Gene Expression Omnibus GSE266291. scRNAseq of DHT treated islets is available at GSE201256. *Pink1*KO kidney bulkRNAseq is available at GSE217775. Patchseq data is available at GSE270484, GSE124742, and GSE164875.

## Code Availability

Code for the analyses reported in this manuscript, including usage of SnakeClassifier and BlackSwanClassifier is available via GitHub at https://github.com/FMJLabTulane/MitochondriaAI while the Mitochondrial Function Index calculator codex workflow is available at https://github.com/FMJLabTulane/MFI.

## Acknowledgements

This work was supported by National Institutes of Health grants DK074970 and P20GM152305 (FM-J), K99DK140067 (MMFQ), DK120447 (PEM), U.S. Department of Veterans Affairs Merit Award BX005812 (FM-J), and the Tulane Center of Excellence in Sex-Based Precision Medicine (FM-J). PEM was supported by Canadian Institutes of Health Research grant 186226, Canada Research Chair in Islet Biology, Alberta-Helmholtz Diabetes Research School, the Alberta Innovates Scholarship in Data-Enabled Innovation, and the Sir Fredrik Banting and Dr. Charles Best Canada Graduate Scholarship (TdS). The preparation of human pancreatic islets provided by the Integrated Islet Distribution Program (IIDP) (RRID:SCR_014387) at City of Hope were funded by NIH grant 2UC4DK098085. This manuscript used data acquired from the Human Pancreas Analysis Program (HPAP-RRID:SCR_016202) Database (https://hpap.pmacs.upenn.edu/), a Human Islet Research Network (RRID:SCR_014393) consortium (UC4-DK-112217, U01-DK-123594, UC4-DK-112232, and U01-DK-123716).

## Contributions

MMFQ analysis, KANN MFI design, dataset curation, manuscript writing and edits. CD and PM-J SnakeClassifier and BlackSwanClassifier design and implementation. CD, PM-J, NH, and FMJ contributed ideas, manuscript writing and finals edits. PEM and TdS provided data for Fig.5m. FMJ supervision.

