## Supplemental_file for "Supervised machine learning identifies impaired mitochondrial quality control in β cells with development of type 2 diabetes"

[1]Tulane Center of Excellence in Sex-based Precision Medicine, New Orleans, LA, USA. [2]Section of Endocrinology, Deming Department of Medicine, Tulane University School of Medicine, New Orleans, LA, USA. [2]Southeast Louisiana VA Medical Center, New Orleans, LA, USA. [4]Algorithme, Paris, Île-de-France, France. [5]Department of Pharmacology, University of Alberta, Edmonton, AB T6G2R3, Canada. [6]Alberta Diabetes Institute, University of Alberta, Edmonton, AB T6G2R3, Canada

Keywords: Type 2 diabetes; Mitophagy; Rule based classifier; Machine learning; Disease prediction

Index:

**Supplementary Figure 1:** Runtime scaling and interpretability

**Supplementary Figure 2:** Threshold behavior and stratified predictions

**Supplementary Figure 3:** Regression analysis between β cell identity pathways mean disease score and HbA1c

**Supplementary Figure 4:** Sex-stratified pathways in β cell disease states

**Supplementary Figure 5:** Stress pathways and electrophysiology link PINK1 status and mitochondrial fitness to β cell function

**Supplementary Figure 6:** Sex and treatment stratified expression of pro-inflammatory genes in β cell subtypes

**Supplementary Figure 7:** Seed robustness and sex-stratified pathway enrichment

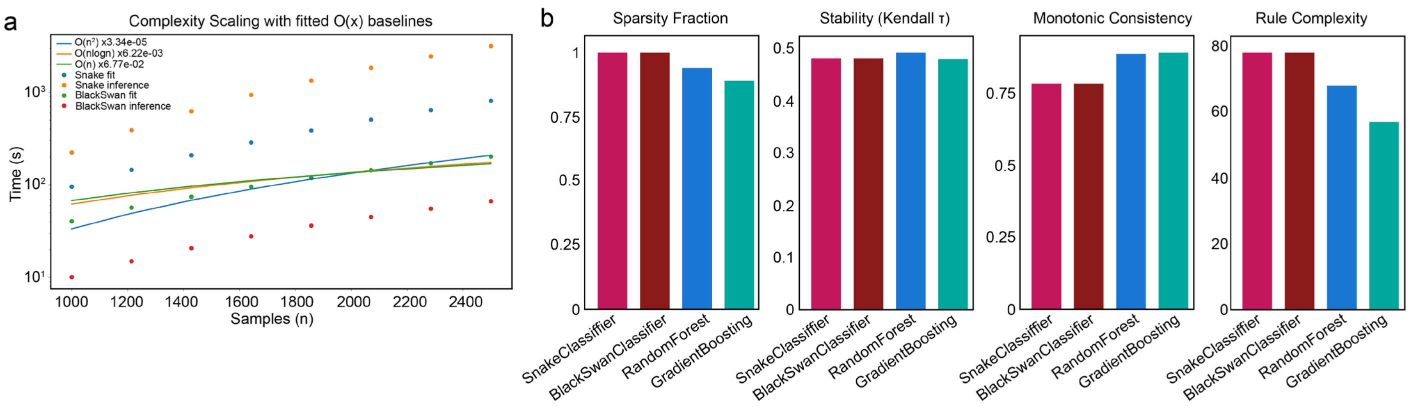


**Supplementary Figure 1 | Runtime scaling and interpretability**

**a,** Training fit and inference times for SnakeClassifier and BlackSwanClassifier for n=1000-2500 cells (log-time), with fitted $O(n^2)$, $O(n\log n)$, and $O(n)$ baselines; both methods scale near $O(n)$-$O(n\log n)$, well below quadratic scales. **b,** Interpretability metrics for SnakeClassifier, BlackSwanClassifier, Random Forest and Gradient Boosting. Sparsity fraction (SnakeClassifier/BlackSwanClassifier = 1; RF = 0.94; GBM = 0.89), stability of feature ranks (Kendall τ = 0.48-0.49), monotonic consistency (SnakeClassifier/BlackSwanClassifier = 0.78; RF = 0.883; GBM = 0.888), and rule complexity (SnakeClassifier/BlackSwanClassifier = 77.6; RF = 67.6; GBM = 56.6).

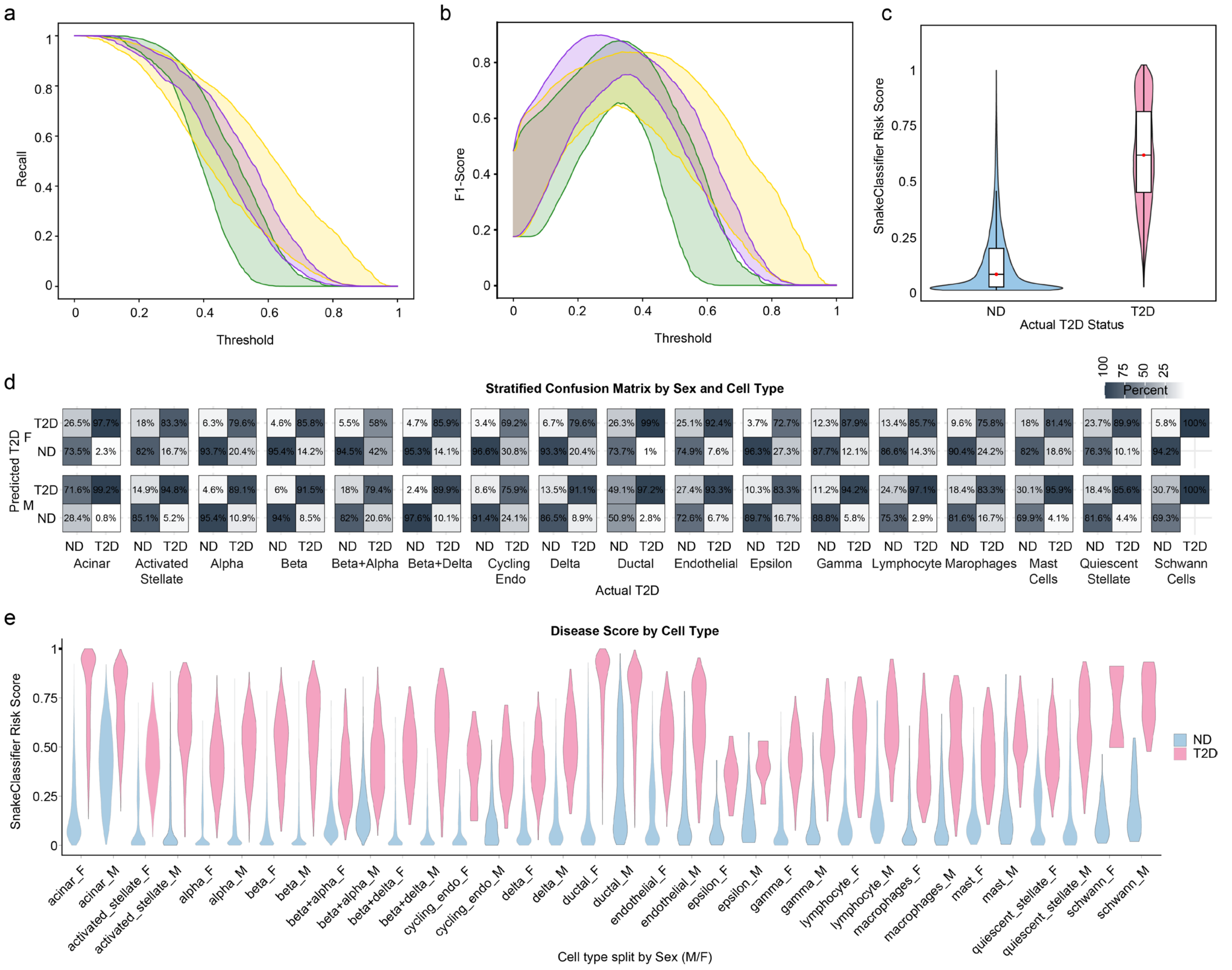


### Supplementary Figure 2 | Threshold behavior and stratified predictions

**a,** Recall as a function of decision threshold for SnakeClassifier, Random Forest and Gradient Boosting; shaded bands indicate variability across resampling. **b,** F1-score versus threshold for the same models, showing the operating range that maximizes balanced accuracy. **c,** Distribution of SnakeClassifier risk scores (0-1, higher = T2D) by true status (ND vs T2D), indicating clear separation. **d,** Stratified confusion matrices by sex and cell type; entries are row-normalized percentages (ND/T2D columns by actual label). **e,** Per-cell disease scores by cell type split by sex (violins; ND blue, T2D pink), highlighting lineage and sex-specific heterogeneity in predicted T2D burden.

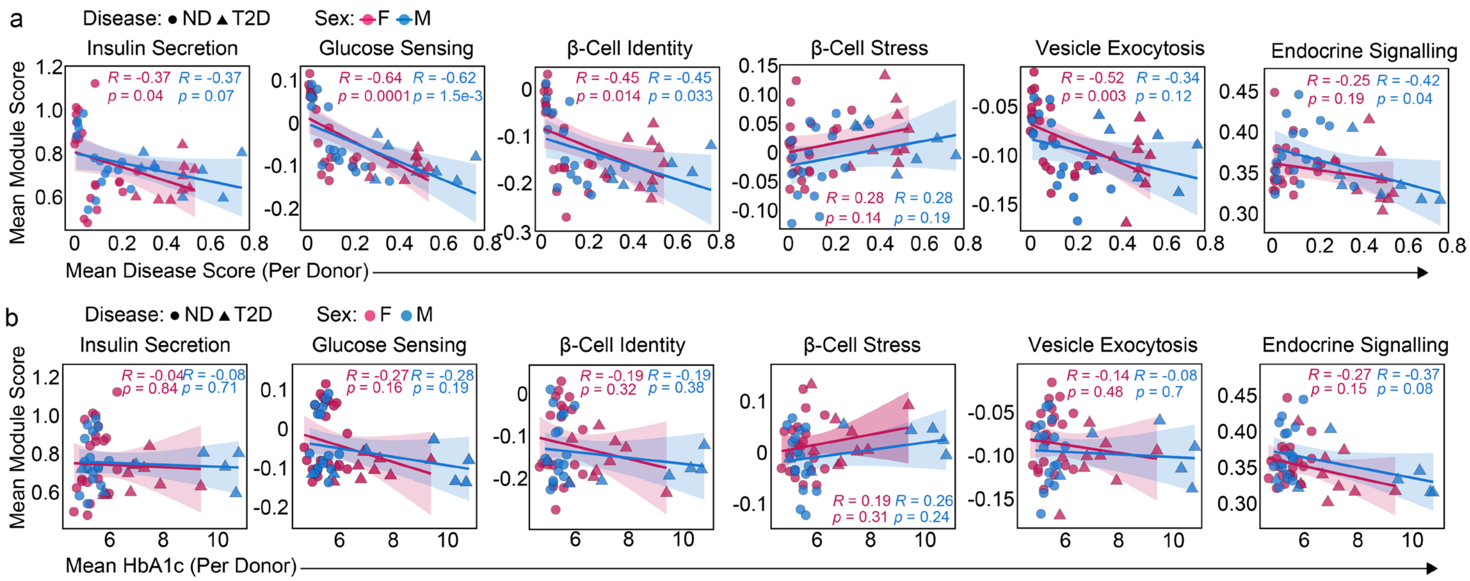


**Supplementary Figure 3 | Regression analysis between β cell identity pathways mean disease score and HbA1c**

**a,** Scatter plots showing the relationship between mean disease risk score and module scores for each gene set, stratified by sex, for ND and T2D donors. Correlation coefficients (R) and FDR-adjusted p-values are shown. **b,** Correlation of mean gene module scores with donor HbA1c (%), stratified by sex and β-cell subtype. Pearson R and FDR-adjusted p-values are indicated for each gene set.

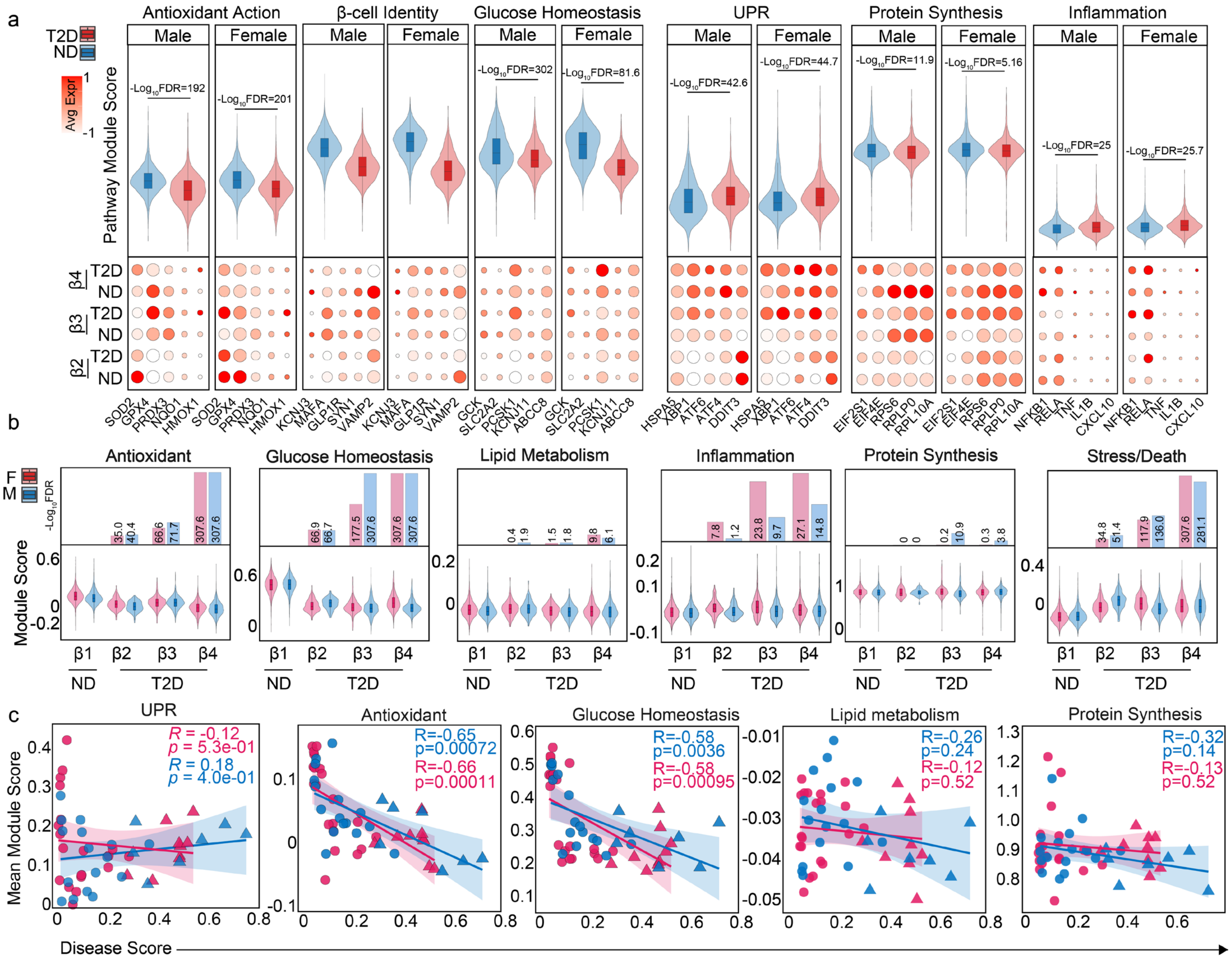


**Supplementary Figure 4 | Sex-stratified pathways in β cell disease states**

**a,** Pathway-module scores (violin plots) for ND vs. T2D, stratified by sex for six programs (antioxidant, β cell identity, glucose metabolism, UPR, protein synthesis, inflammation). Beneath each panel representative gene expression dotplots summarize program genes; dot size denotes fraction of expressing cells and fill intensity indicates average expression (scaled). **b,** Module scores within β cell subtypes (β1-2), split by disease and sex (violin plots); insets report -log$_{10}$(FDR) for T2D/ND contrasts by sex within each cluster. **c,** Association between module scores and SnakeClassifier disease score for β cell groups; lines show sex-specific linear fits with 95% CI, Pearson R and p-value are annotated.

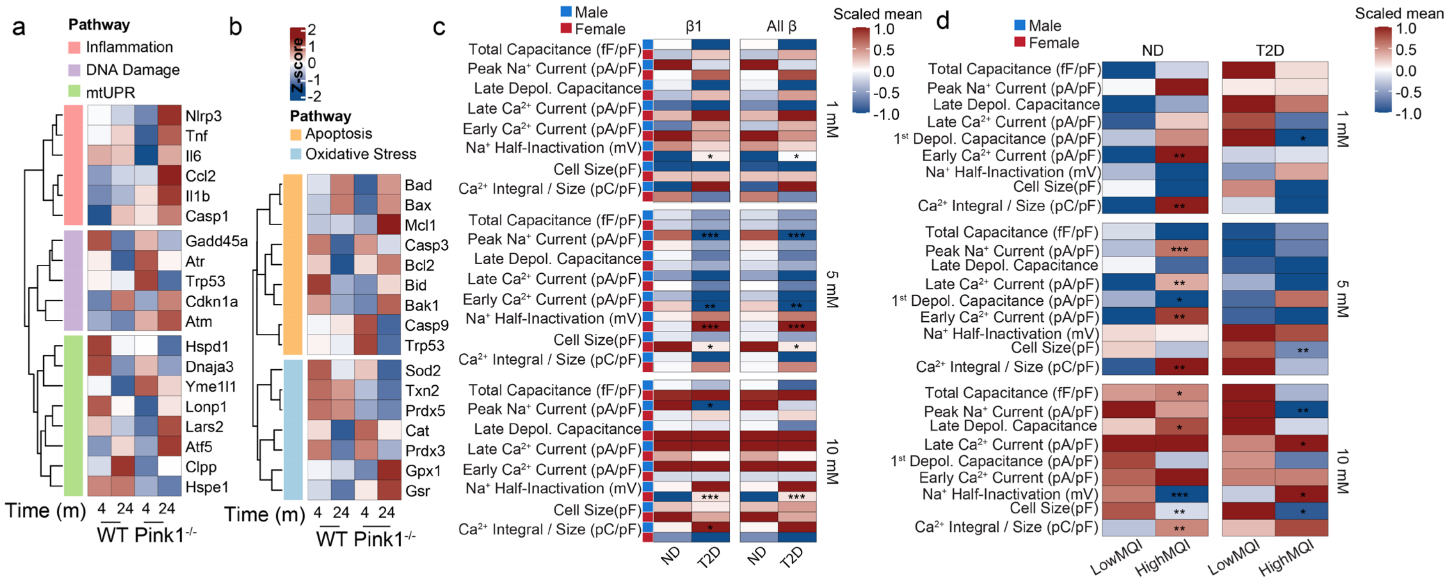


**Supplementary Figure 5 | Stress pathways and electrophysiology link *PINK1* status and mitochondrial fitness to β cell function**

**a,** Heatmap of inflammation, DNA-damage, and mitochondrial unfolded-protein response (mtUPR) genes showing scaled mean expression in WT and *Pink1*$^{-/-}$ islets at 4 and 24 months; hierarchical ordering highlights coordinated induction of innate-immune (e.g., *Nlrp3*, *Tnf*, *Il6*) and mtUPR chaperones. **b,** Companion heatmap for apoptosis and oxidative-stress programs (e.g., *Bax*, *Casp3*, *Sod2*, *Gpx1*), indicating progressive stress signaling in *Pink1*$^{-/-}$ relative to WT with age. **c,** Patch-seq electrophysiology summary matrices for β1 cells and all β-cells (ND and T2D), comparing *PINK1*$^{-}$ vs *PINK1*$^{+}$ states against stimulus steps (1, 5, 10 mM). Entries display scaled means for capacitance, $Na^{+}$/$Ca^{2+}$ currents (peak/late/early), $Na^{+}$ half-inactivation, cell size, and $Ca^{2+}$ integral/size; asterisks denote significant differences (two-sided tests, FDR-adjusted). **d,** Electrophysiology stratified by mitochondrial fitness group (Low-MFI vs High-MFI) within ND and T2D against the same stimulus steps. Patterns recapitulate altered excitability and calcium handling in low-fitness and *PINK1*$^{-}$ states. All p-values are two-sided with Benjamini–Hochberg FDR control as detailed in Methods.

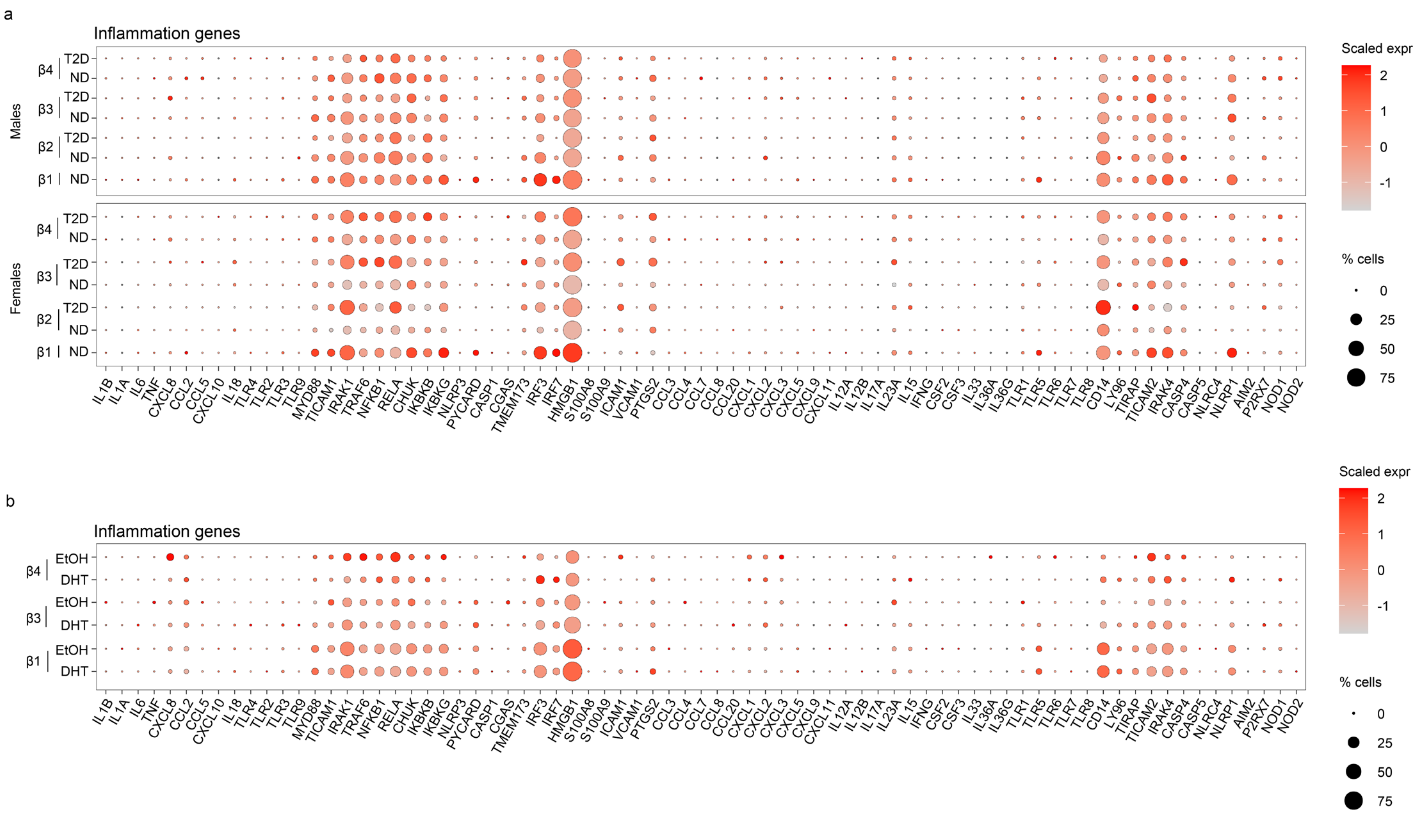


**Supplementary Figure 6 | Sex and treatment stratified expression of pro-inflammatory genes in β cell subtypes**

**a,** Dotplots for a curated inflammation panel for β1-4, split by sex and disease. Color encodes scaled average expression; dot size encodes the fraction of expressing cells. Within each sex, rows are ordered β1-4 with ND preceding T2D. **b,** Dotplots for a curated inflammation within male β cells under DHT or vehicle (EtOH); rows are β1-4 with DHT preceding EtOH. Color encodes scaled average expression; dot size encodes the fraction of expressing cells.

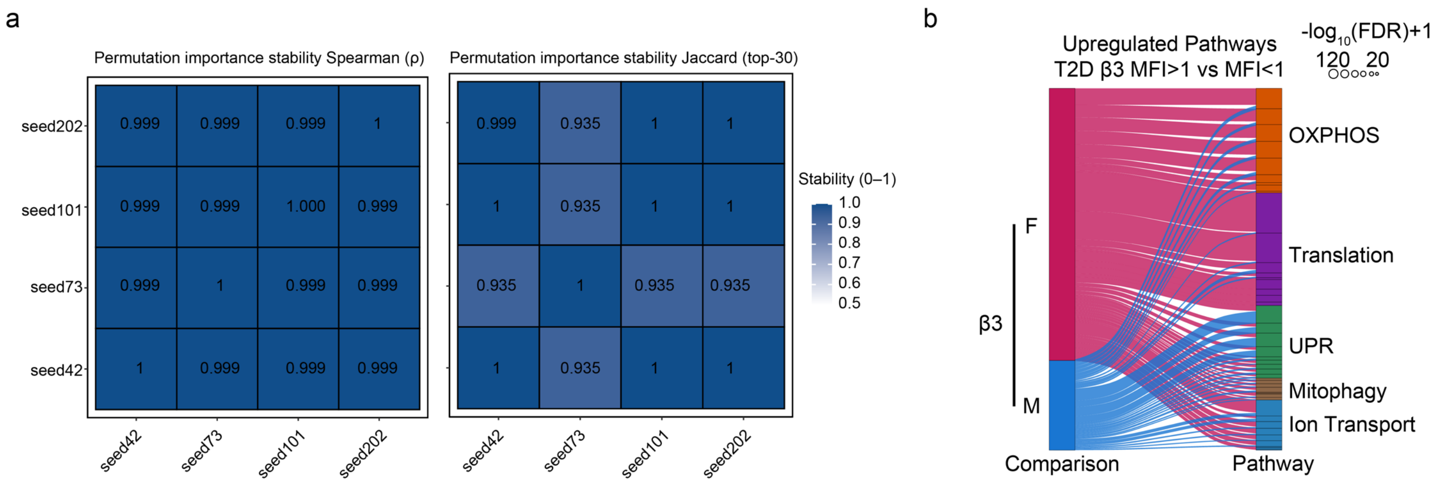


**Supplementary Figure 7 | Seed robustness and sex-stratified pathway enrichment**

**a,** Stability of permutation importance for random seeds (42, 73, 101, 202). Left: Spearman rank correlations of full importance vectors (ρ = 0.999-1). Right: Jaccard similarity of the top30 features (0.935-1). **b,** Pathways upregulated in T2D β3 cells with high mitochondrial function index (MFI>1) versus low (MFI<1), stratified by sex. Sankey ribbons connect the comparison group to pathway families (e.g., OXPHOS, Translation, UPR, Mitophagy, Ion transport); ribbon width encodes significance (-$\log_{10}$(FDR)+1).
